# VEGF-C promotes brain-derived fluid drainage, confers neuroprotection, and improves stroke outcomes

**DOI:** 10.1101/2023.05.30.542708

**Authors:** Ligia Simoes Braga Boisserand, Jean Bouchart, Luiz Henrique Geraldo, Seyoung Lee, Basavaraju G. Sanganahalli, Maxime Parent, Shenqi Zhang, Yuechuan Xue, Mario Skarica, Justine Guegan, Mingfeng Li, Xiodan Liu, Mathilde Poulet, Michael Askanase, Artem Osherov, Myriam Spajer, Marie-Renee El Kamouh, Anne Eichmann, Kari Alitalo, Jiangbing Zhou, Nenad Sestan, Lauren H. Sansing, Helene Benveniste, Fahmeed Hyder, Jean-Leon Thomas

## Abstract

Meningeal lymphatic vessels promote tissue clearance and immune surveillance in the central nervous system (CNS). Vascular endothelium growth factor-C (VEGF-C) is essential for meningeal lymphatic development and maintenance and has therapeutic potential for treating neurological disorders, including ischemic stroke. We have investigated the effects of VEGF-C overexpression on brain fluid drainage, single cell transcriptome in the brain, and stroke outcomes in adult mice. Intra-cerebrospinal fluid administration of an adeno-associated virus expressing VEGF-C (AAV-VEGF-C) increases the CNS lymphatic network. Post-contrast T1 mapping of the head and neck showed that deep cervical lymph node size and drainage of CNS-derived fluids were increased. Single nuclei RNA sequencing revealed a neuro-supportive role of VEGF-C via upregulation of calcium and brain-derived neurotrophic factor (BDNF) signaling pathways in brain cells. In a mouse model of ischemic stroke, AAV-VEGF-C pretreatment reduced stroke injury and ameliorated motor performances in the subacute stage. AAV-VEGF-C thus promotes CNS-derived fluid and solute drainage, confers neuroprotection, and reduces ischemic stroke damage.

**Short abstract:** Intrathecal delivery of VEGF-C increases the lymphatic drainage of brain-derived fluids confers neuroprotection, and improves neurological outcomes after ischemic stroke.

---

Lymphatic vessels are present in tissues of most body organs where they clear metabolic waste and control the immune surveillance (Aspelund *et al*., 2015; Oliver *et al*., 2020). In contrast, brain and spinal cord tissues are devoid of lymphatics, despite their intense metabolic activity. A large body of studies conducted in rodents over the last decade indicates that the clearance of solute waste from CNS tissues involves a unique synergy between the cerebrospinal fluid (CSF), the glymphatic system, and the lymphatic system connecting the dura mater to the CNS-draining lymph nodes (Proulx, 2021; Rustenhoven *et al*., 2021). The regulation of meningeal and peri cranial lymphatic drainage is thus of prime importance for maintaining brain tissue homeostasis, especially in neuropathological conditions, for example, when interstitial fluid (ISF) and waste accumulate after an acute brain injury.

CSF is continuously generated by the choroid plexus and circulates through the CNS internal ventricles, the subarachnoid space, and cisterns, as well as along the perivascular spaces of all cerebral vessels (Abbott *et al*., 2018). At the level of the perivascular conduits, CSF exchanges with the ISF of the neuropil through the glymphatic system, thereby facilitating waste clearance from the CNS (Iliff *et al*., 2012; Plog and Nedergaard, 2018; Benveniste *et al*., 2019). This glymphatic system thus generates an outflow of CNS-derived fluids and waste solutes (CSF/ISF) that subsequently drains out of the skull and the vertebral canal.

In mice, CNS-derived fluids exit the skull and the vertebral column through the cribriform plate and other perineural cranial and vertebral routes and then drain via afferent lymphatics into lymph nodes (LNs) located in the mandibular, cervical, and lumbosacral regions (Laman and Weller, 2013; Ma *et al*., 2019; Norwood *et al*., 2019; Proulx, 2021). A body of recent studies also indicates that brain-derived waste solutes and antigens are collected by meningeal lymphatic vessels (MLVs) positioned in the dura mater (Aspelund *et al*., 2015; Da Mesquita *et al*., 2018; Louveau *et al*., 2018; Ahn *et al*., 2019). MLVs are also present in human and nonhuman primates (Absinta *et al*., 2017), and they extend around the brain, spinal cord, and sensory nerve ganglia (Antila *et al*., 2017; Louveau *et al*., 2018; Jacob *et al*., 2019). Drainage from the MLVs to the afferent lymphatics connecting with the cervical LN has been demonstrated using real-time imaging (Ahn *et al*., 2019), but there is a gap in knowledge as to how stimulation of lymphatic function modulates solute drainage from the CNS in living mice.

Gain or loss of MLVs can be induced by a variety of surgical, pharmacological, or gene transfer approaches, which have been used in combination with different mouse models of chronic brain disorders such as Alzheimer’s disease and multiple sclerosis (Antila *et al*., 2017; Da Mesquita *et al*., 2018; Louveau *et al*., 2018). Collectively, these studies revealed that dysfunctional lymphatic drainage may play a role in the development of neurodegenerative diseases. Moreover, the MLVs regulate immune responses in the CNS by transporting CNS-derived antigens to the cervical LNs and enabling T cell priming, as shown in brain tumor cell inoculation (Hu *et al*., 2020; Song *et al*., 2020). Finally, in acute brain injury, lymphatic disruption has been associated with worsened outcomes (Yanev *et al*., 2020). MLVs have been reported to regulate the extent of edema, the degree of microglial activation, and overall neuronal degeneration, in a brain concussion model (Bolte *et al*., 2020) and in experimental stroke experiments conducted in the zebrafish (Chen *et al*., 2019) and mice (Yanev *et al*., 2020). These findings suggest that the expansion of MLVs and their downstream lymphatic vasculature may enhance the upstream drainage of CNS-derived fluids, waste, and immune cells following an ischemic stroke injury, thereby reducing ischemic edema as well as accumulation of debris and inflammatory cells, with improved of outcomes after stroke. Lymphatic development and growth is controlled by VEGF-C, which activates the VEGFR3 tyrosine kinase receptor on the surface of lymphatic endothelial cells (Lohela *et al*., 2009). We and others have shown that the VEGF-C/VEGFR3 signaling system controls the growth and plasticity of meningeal lymphatics and that intra-CSF delivery of VEGF-C, either as a recombinant protein or via AAV-VEGF-C, stimulated MLV growth (Aspelund *et al*., 2015; Louveau *et al*., 2015; Antila *et al*., 2017; Song *et al*., 2020). Furthermore, VEGF-C activates brain neural stem/progenitor cells (Le Bras *et al*., 2006; Calvo *et al*., 2011; Hayakawa *et al*., 2011; Han *et al*., 2015), which may induce protective and repair processes in CNS tissues after traumatic or ischemic injuries.

Here, we assessed the consequences of intrathecal delivery of AAV-VEGF-C on CSF and solute drainage as well as gene expression in the brain. Using contrast-enhanced magnetic resonance imaging (MRI) for real-time tracking of a CSF-injected gadolinium-based solute tracer, we found that AAV-VEGF-C delivery into the CSF did not alter glymphatic drainage and CSF efflux through the cribriform plate but increased the volume and drainage of deep cervical lymph nodes (dCLNs). Single nuclei RNA-sequencing (snRNA-seq) analysis of cerebral cells showed that intrathecal treatment with AAV-VEGF-C upregulated a network of pathways associated with calcium and BDNF signaling in resident brain cells, including subsets of interneurons, astrocytes, and endothelial cells. We next investigated whether CSF delivery of AAV-VEGF-C or recombinant VEGF-C would improve the anatomical and functional outcomes after an acute brain injury, using the clinically relevant mouse model of focal ischemic stroke induced by transient middle cerebral artery occlusion (tMCAO). We found that AAV-VEGF-C pretreatment, but not post-stroke treatment with rVEGF-C, reduces the stroke lesion volume and peri-lesional inflammation while improving motor behavior. This study shows that intrathecal delivery of AAV-VEGF-C simultaneously improves CSF lymphatic drainage, increases myelin gene expression, and promotes neurosupportive signals in brain neural cells, thereby potentially counteracting the deleterious effects of ischemic injury.

## Results

### VEGF-C pretreatment promotes brain-derived fluid drainage

To assess the consequences of intra-CSF AAV-VEGF-C delivery on CSF/ISF outflow drainage via the glymphatic system *in vivo*, we conducted quantitative magnetic resonance imaging (MRI). Mice received an intra-cisterna magna (ICM) injection of AAV-VEGF-C or AAV-Control (CTRL) (Fig.1A). Four weeks after treatment, we performed MRI using variable flip angle 3D spoiled gradient echo (VFA-SPGR) T1 mapping in combination with ICM administration of gadoteric acid (Gd-DOTA, molecular weight 558 Da) (Koundal *et al*., 2019; Xue *et al*., 2020). All MRI was carried out on a 9.4T MRI instrument on ketamine/xylazine anesthetized mice (*n* = 10 mice per group). Quantitative analysis of the T1 maps acquired ∼1hr after CSF administration showed that brain-wide glymphatic transport of Gd-DOTA was unaffected by VEGF-C treatment compared to controls (AAV-CTRL: 47,567 ± 3,095 voxels and AAV-VEGF-C: 49,376 ± 2,477 voxels, *P =* 0.65, Unpaired t-test). As shown in Fig. 1B, brain-wide glymphatic transport is visualized as a 3D-volume rendered binary map in control (blue), and VEGF-C treated (green) mice. Specifically, the binary maps capture all voxels with T1 values in the range of 1-1700 ms which represents brain tissues with the uptake of Gd-DOTA as a measure of glymphatic transport (Xue *et al*., 2020). In the same mice, we also evaluated drainage of Gd-DOTA via cranial outflow pathways. Quantification of Gd-DOTA efflux to nasal cavity via the cribriform plate in the olfactory region did not reveal any differences between control and VEGF-C treated mice (AAV-CTRL: 1,690 ±160 voxels and AAV-VEGF-C: 1,908 ± 127 voxels, *P =* 0.30, Unpaired t-test) (Fig. 1C).

**Figure 1.**
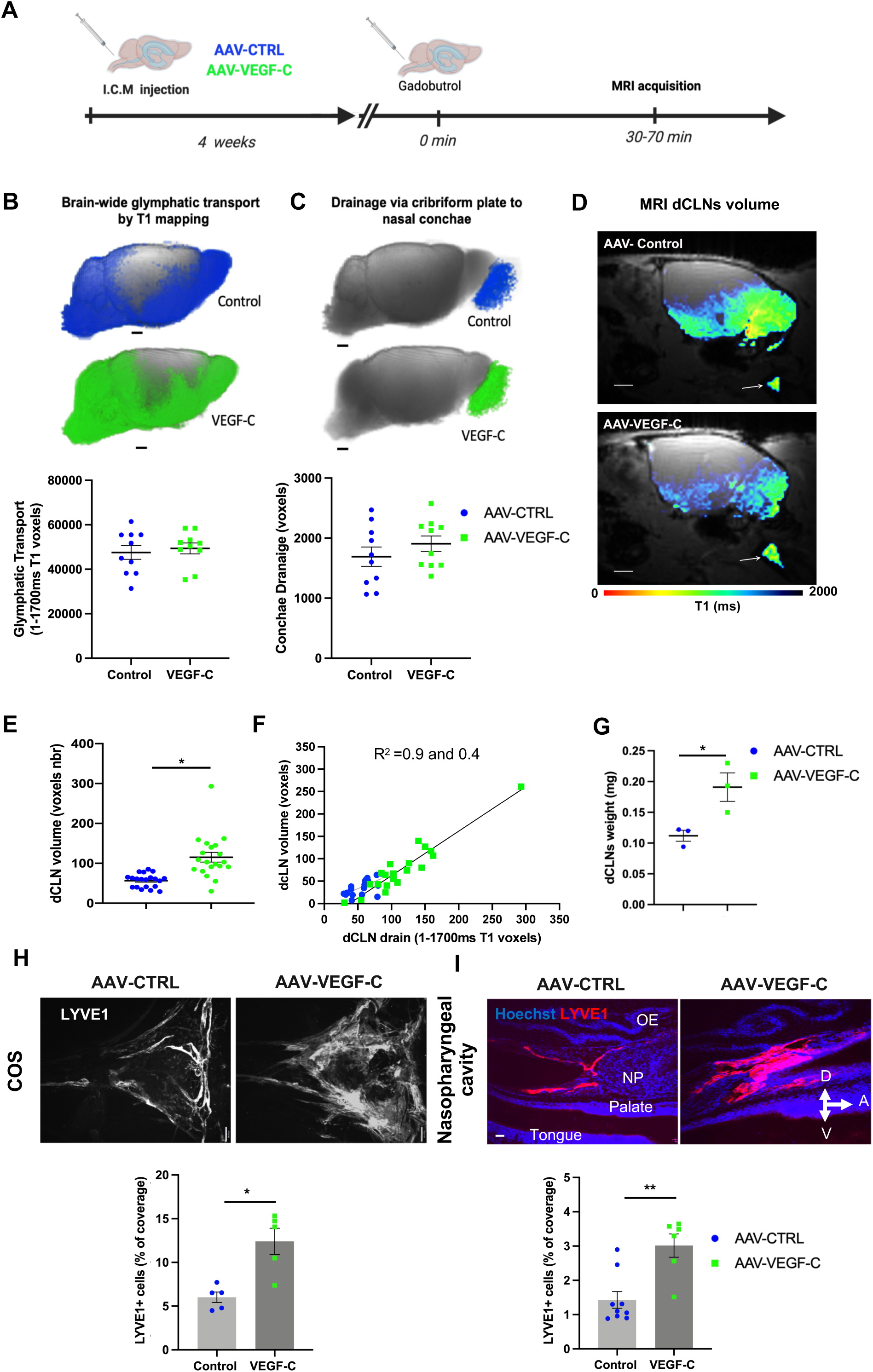
AAV-VEGF-C promotes drainage of CSF-injected Gd-DOTA into dCLNs. **(A)** Schematic diagram of the experimental procedure. Eight weeks old (8 wo) mice were analyzed by DCE-MRI after a prior ICM injection of either AAV-VEGF-C or AAV-control at 4 wo (*n* = 10/group). After intra-CSF administration of Gd-DOTA, T1 mapping technique was applied to detect the **(B)** glymphatic transport in the normal brain tissue, which is characterized by T1 values in the 1-1700ms range. The number of voxels was comparable between the two groups of mice, indicating that the glymphatic transport was not altered by VEGF-C preconditioning (*P=0.65,* Unpaired t-test). **(C)** Using the same approach, we evaluated the extracranial outflow of Gd-DOTA efflux in the cribriform plate-olfactory epithelium area. The two groups of mice showed a similar outflow of Gd-DOTA efflux (*P=0.30,* Unpaired t-test). Anatomical evaluation of dcLNs showing that the nodal volume and the intra-nodal circulation of Gd-DOTA were both increased in VEGF-C preconditioned mice compared to controls. **(D)** Plane view sagittal to the brain at the level of the dCLN showing the T1 signal mapped to the node (white arrows) and an enlarged dCLN in the AAV-VEGF-C mouse compared to CTRL. Scale bar: 1.5mm. **(E)** Measurement of dcLNs volume as several Gd-DOTA-positive voxels (AAV-VEGF-C: 115.2 ± 12.1 and AAV-control: 56.5 ± 3.7, \*\*\**P* < *0.0001,* Mann-Whitney test). A linear regression analysis showed that the dcLN volume is directly related to the intra-nodal circulation of Gd-DOTA (AAV-VEGF-C: R^2^ = 0.89 \*\*\**P < 0.0001;* AAV-control: R^2^ = 0.39 \*\**P* < 0.004) **(F)**. Postmortem analysis showing that dcLN weight was increased in VEGF-C-preconditioned mice (AAV-VEGF-C: 0.19 ± 0.02 and AAV-control: 0.11 ± 0.01, \**P* < *0.03, n =* 9 dcLNs (3 dcLNs/each measurement), (Unpaired t-test) **(G)**. MLV growth is increased in VEGF-C-preconditioned mice. The surface of LYVE1^+^ labeling (%) was measured at the confluence of sinuses (*n* = 5 mice/group, \**P* < 0.05, Mann-Whitney test) (**H**). Representative images of extracranial lymphatics in the nasopharyngeal area (*n* = 6-9 mice/group, \*\**P* < 0.005, Mann-Whitney test). Data are represented as mean ± S.E.M (**I**). A: anterior, D: dorsal, V: ventral. NP: nasopharynx, OE: olfactory epithelium. Scale bars: Scale bars: 1.5mm (**D**), 100μm (**H**), 35μm (**I**).

Next, we evaluated the drainage of Gd-DOTA to dCLNs, using a quantitative T1 map analysis of Gd-DOTA uptake (Fig. 1D). We first used anatomical MRI to measure the volume of the left and right dCLNs on the corresponding anatomical MRI. In the subsequent analyses, each left and right dCLN was treated as an independent measure as there was no difference between left and right dCLN volume in control mice (Left 0.35 ± 0.09 mm^3^ versus Right 0.32 ± 0.11 mm^3^, *P* = 0.434). Interestingly, the comparison of dCLN volume across groups revealed that dCLN volume was significantly increased in VEGF-C pretreated mice when compared to controls (dCLN volume AAV controls: 0.34 ± 0.10 mm^3^ versus dCLN volume AAV-VEGF-C: 0.69 ± 0.32 mm^3^, \*\*\*\**P* < 0.0001). Furthermore, we found a statistically significant increase in voxel volume as defined by T1 values in VEGF-C-pretreated mice compared to controls, signifying overall more drainage (Fig. 1E). A linear regression analysis with VEGF-C treatment showed that dCLN volume was directly related to Gd-DOTA uptake into the dCLNs, suggesting that VEGF-C-induced enlargement of dCLNs is promoting drainage of CSF from the meningeal lymphatic and the glymphatic systems (Fig. 1F). Finally, postmortem analysis further validated that the VEGF-C pre-treatment enhances MLVs hence it increases the weight of dCLNs (Fig. 1G).

Following MRI, mice were euthanized to assess MLV morphology along the dura mater-to-cervical LN drainage pathway. As expected, Intra-cisternal delivery of VEGF-C was found to increase the coverage and diameter of meningeal lymphatics as expected (Fig. 1H), but it also remodeled extracranial nasopharyngeal lymphatics (Fig. 1I). In contrast, the morphology of lymphatics in the ear skin was not altered by ICM injection of AAV-VEGF-C (Supplementary Fig. 1A, B), indicating that intra-CSF VEGF-C acts locally on lymphatics of the CSF drainage pathway, but not on extracranial peripheral lymphatics of the head.

Therefore, intra-CSF VEGF-C expands the meningeal, extracranial, and LN compartments of the dura mater-to-LNs pathway. This suggests that VEGF-C promotes the drainage of CNS-derived fluids to their collecting LNs via enlargement of MLVs but not glymphatic transport.

### snRNA-seq analysis of forebrain cells after intra-CSF AAV-VEGF-C delivery

To determine if and how AAV-VEGF-C delivery into the CSF impacts the CNS parenchyma, we performed immunostaining and snRNA-seq of the cortex and striatum tissues of the forebrain. Following ICM injection of a control AAV serotype 9-GFP vector, we were able to identify AAV_9_-infected brain cells by fluorescence. GFP was detected in all different brain regions among subsets of perivascular smooth muscle cells, NeuN^+^ neurons, and Olig2^+^ oligodendroglial cells, but only in a few endothelial cells and astrocytes (Supplementary Fig. 1C-J). Using AAV-VEGF-C-treated mice, we then assessed the pro-neurogenic effect of VEGF-C previously reported in the subventricular zone along lateral ventricles (Calvo *et al*., 2011; Han *et al*., 2015). The number of subventricular doublecortin (DCX^+^) neuroblasts was increased in AAV-VEGF-C- pretreated mice compared to controls as early as 7 days after ICM injection of AAVs (Supplementary Fig. 1K-N). The ICM administration of AAV-VEGF-C thus triggers subventricular neurogenesis like the previously reported intracerebral injection of recombinant VEGF-C (Han *et al*., 2015). Altogether these results suggests that beyond a specific effect on neural stem/progenitor cells, CSF delivery of AAV-VEGF-C shall affect other cell types and processes within the brain.

We used snRNA-seq to further assess the transcriptional changes induced by AAV-VEGF-C pretreatment in brain cells. SnRNA-Seq is well suited for the identification of cellular heterogeneity in brain tissues as well as gene expression changes in brain cell subpopulations between different experimental groups (Zeisel *et al*., 2015; Schirmer *et al*., 2019). SnRNA-seq profiling was performed on cortical and striatal areas dissected from AAV-VEGF-C and AAV-CTRL mice (*n* = 5 mice/group) (Fig. 2A). We sequenced 78,728 nuclei, with a median of 1,627 genes and 2,550 transcripts per nucleus (Supplementary Table 1, Supplementary Fig. 2). We annotated 18 main cell clusters (Fig. 2B). Expression of lineage marker gene signatures characterized in previous snRNA-seq studies was used to identify *Syt1* (Synaptotagmin) for neurons, *Aqp4* (aquaporin 4) for astrocytes, *Tmem119* (transmembrane protein 119) for microglia, *Mbp* (myelin basic protein) for oligodendrocytes, *Pdgfr*α (platelet-derived growth factor receptor alpha) for oligodendrocytes precursor cells (OPCs) and *Cldn5* (claudin 5) for endothelial cells (He *et al*., 2018; Schirmer *et al*., 2019) (Fig. 2C, Fig. 2D, Supplementary Figs. 3, 4). As expected, *Cux2*-expressing cortical pyramidal neurons and *Spink8*-expressing hippocampal pyramidal neurons were lacking in striatal samples, *Sst*- and *Vip*-expressing inhibitory interneurons concentrated in cortical samples, and neurons expressing *Sv2c* (synaptic vesicle glycoprotein 2C) were predominant in the striatum (Dardou *et al*., 2011), while glial cells, as well as vascular cells, were equally distributed between cortex and striatum, which supports the accuracy of the clustering (Supplementary Figs. 5A, C; Supplementary Table 2). We found similar numbers of neuronal, glial, and non-neural cells between AAV-Ct and AAV-VEGF-C pretreated mice (Supplementary Figs. 5B, C; Supplementary Table 3). The tSNE representation showed no major changes in the clustering after AAV-VEGF-C treatment (Supplementary Fig. 5C). Alteration of transcriptional expression was, however, detectable in several clusters such as *Sv2c*- and inhibitory neurons, meningeal fibroblasts, and smooth muscle cells (Supplementary Fig. 2B). To gain insight into the cell-type changes in gene expression between experimental conditions, we performed a differential expression analysis within the different clusters of the dataset.

**Figure 2.**
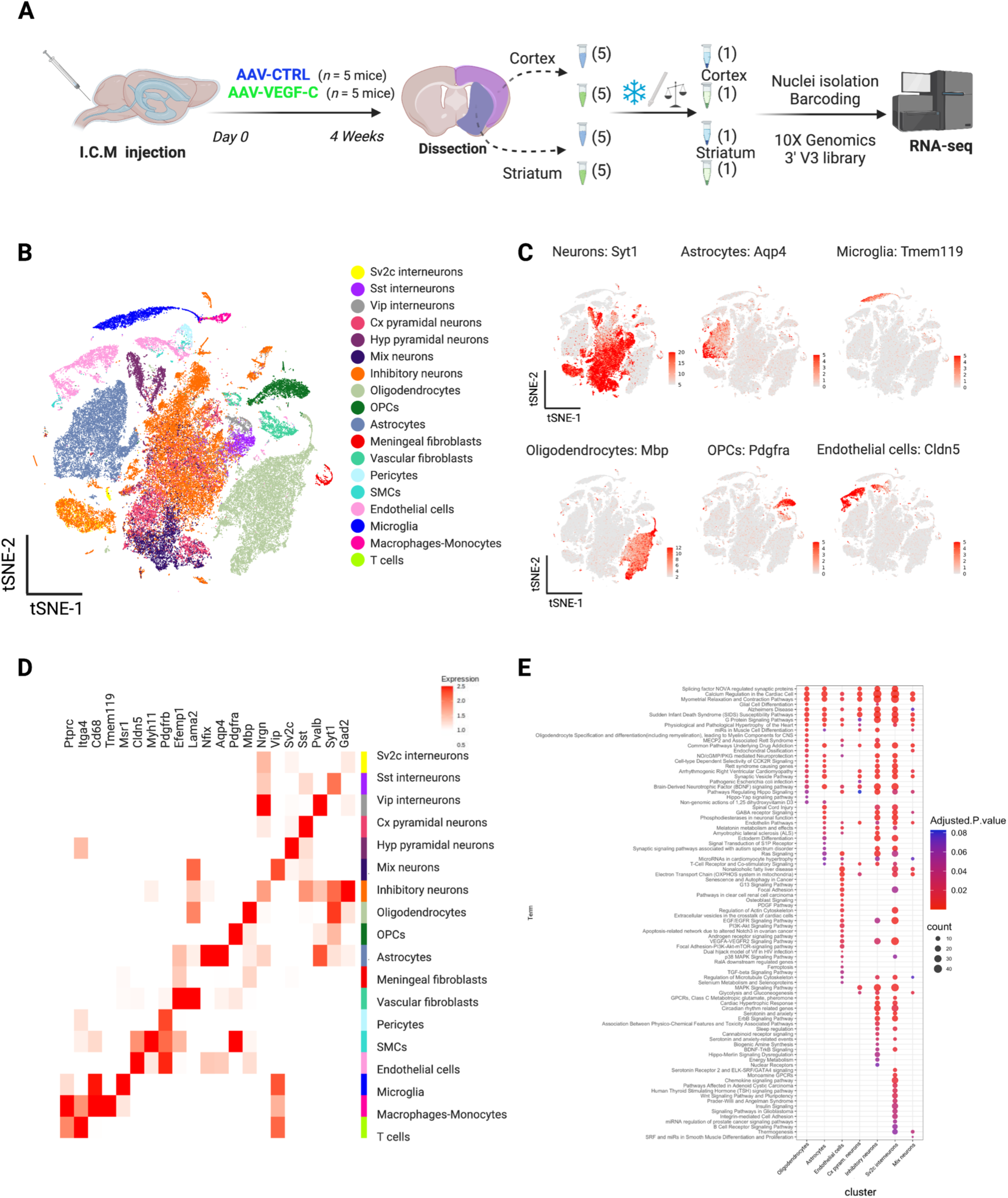
snRNA-seq analysis of forebrain cells after intra-CSF AAV-VEGF-C delivery. **(A)** Schematic diagram of the experimental procedure. **(B)** Cell-type specific clustering for the whole sample contribution to each cluster. **(C)** tSNE plot highlighting markers genes expression for neurons, astrocytes, microglia, oligodendrocytes, OPCs, and endothelial cells. **(D)** Expression heatmap of the specific marker genes expression for each cell cluster. **(E)** WikiPathways analysis shows the signaling pathways alterations induced by VEGF-C among brain cell clusters. Pathway enrichment analysis in oligodendrocytes, astrocytes, endothelial cells, cortical pyramidal (cx pyram) neurons), inhibitory neurons, *Sv2c*-interneurons, and Mix neurons. These clusters show more than 180 differentially expressed genes (padj < 0.05 et abs(log2FC) > 0.58) with pathways with at least 20 genes and adjusted p-values < 0.05 in at least 1 cluster.

### Single brain nucleus transcriptome analysis of VEGF-C-responsive gene networks

We first investigated the signaling pathway alterations induced by VEGF-C among brain cell clusters. A pathway enrichment analysis was conducted with the WikiPathways platform using the enrichR package. We focused on clusters showing more than 180 differentially expressed genes (padj < 0.05 et abs(log2FC) > 0.58) and only considered pathways with at least 20 genes and adjusted p-values < 0.05 in at least one cluster for further analysis. The clusters selected for pathway enrichment analysis included endothelial cells, subpopulations of inhibitory neurons (*GAD*^+^), *Sv2c*-interneurons, cortical (cx) pyramidal neurons, endothelial cells, astrocytes, and oligodendrocytes (Fig. 2E). Based on the number of altered expression pathways (*n* > 25), we observed that the clusters most significantly affected by VEGF-C treatment were *Sv2c*-interneurons, inhibitory neurons, as well as endothelial cells, and astrocytes. Pathways of calcium regulation, G protein signaling, Alzheimer’s disease signaling, and BDNF signaling were upregulated in all four clusters. Especially, activation of the BDNF pathway was higher in inhibitory neurons (*n* = 26; adjusted p-value = 2.7.10^-7^) and *Sv2c*-expressing interneurons (*n* = 29; adjusted p-value = 9.02.10^-6^) than in astrocytes (*n* = 12; adjusted p-value = 4.4.10^-5^) and endothelial cells (*n* = 6; adjusted p-value = 0.02). Neurons share with endothelial cells the upregulation of tyrosine kinase receptor signaling pathways, including mitogen-activated protein kinase (MAPK), Vascular Endothelial Growth Factor (VEGF), and Epidermal Growth Factor (EGF). Neurons also showed upregulated pathways in common with astrocytes: phosphodiesterase signaling, serotonin signaling (sudden infant death syndrome), and the regulation of synapse protein synthesis (NOVA-regulated synaptic proteins). Interestingly, signaling pathways of glial cell differentiation and myelination were upregulated together with the BDNF pathway in oligodendrocytes (Fig. 2E).

We then examined specific gene upregulation in the cells of interest. In the brain, calcium is mandatory for the control of synaptic activity and memory formation (Ghosh and Greenberg, 1995). Analysis of the top-upregulated genes indicated that both subsets of neurons and astrocytes increased calcium signaling upon CSF delivery of AAV-VEGF-C. Inhibitory neurons (Fig. 3A) showed overexpression of *Pcp4* (Purkinje cell protein 4), a modulator of calcium binding by calmodulin (Sköld *et al*., 2006, p. 19) (log fold change (logFC): 7.17, adjusted p-value = 3.55E-167), while *Sv2c* interneurons (Fig. 3B) displayed highly upregulated expression of *ATP2B2* (ATPase plasma membrane Ca^2+^ transporting 2a, logFC: 8.56, adjusted p-value: 1.51E-23), a modulator of presynaptic Ca^2+^ homeostasis and H^+^ gradient in synaptic vesicles (Ono *et al*., 2019), and *GRIN2B* (glutamate ionotropic receptor NMDA Type Subunit 2B, logFC: 4.60, adjusted p-value = 1.95E-24), a key regulator of the neuronal intracellular calcium homeostasis (Liu *et al*., 2022) (*ATP2B2*, logFC: 8.56, adjusted p-value: 1.51E-23; *GRIN2B*, logFC: 4.60, adjusted p-value = 1.95E-24). In astrocytes (Fig. 3C), top-upregulated genes included *Calm2* (Calmodulin 2; logFC: 1.87, adjusted p-value = 3.00E-57) and *Nrgn* (Neurogranin; logFC: 9.50, adjusted p-value = 3.70E-47), a postsynaptic calmodulin-interacting protein (Pak *et al*., 2000). These data suggest a potential positive effect of VEGF-C on inhibitory neurons activity and synaptic plasticity. Other top-upregulated genes induced by VEGF-C in brain cells included ferritin heavy chain 1 (*FTH1*) which limits intracellular oxidative stress and production of iron-related protein (Fang *et al*., 2021) (Figs. 3A, D), and the *Slc1a2* solute carrier gene encoding glutamate transporter-1 (GLT-1), the principal transporter of the excitatory neurotransmitter glutamate (Fig. 3D). *FTH1*was overexpressed by inhibitory interneurons (logFC: 27.71, adjusted p-value = 1.96E-256) and endothelial cells (logFC: 70.56, adjusted p-value = 2.21E-31), while *Slc1a2* expression was highly upregulated in endothelial cells (logFC: 17.55, adjusted p-value = 1.04E-31). Overexpression of these genes induced by VEGF-C may mediate neuroprotective effects as *FTH1* limits ferroptosis (Li *et al*., 2021), while *Slc1a2* is expected to prevent glutamate overflow, thereby excitotoxicity, in brain tissues (Harvey *et al*., 2011).

**Figure 3.**
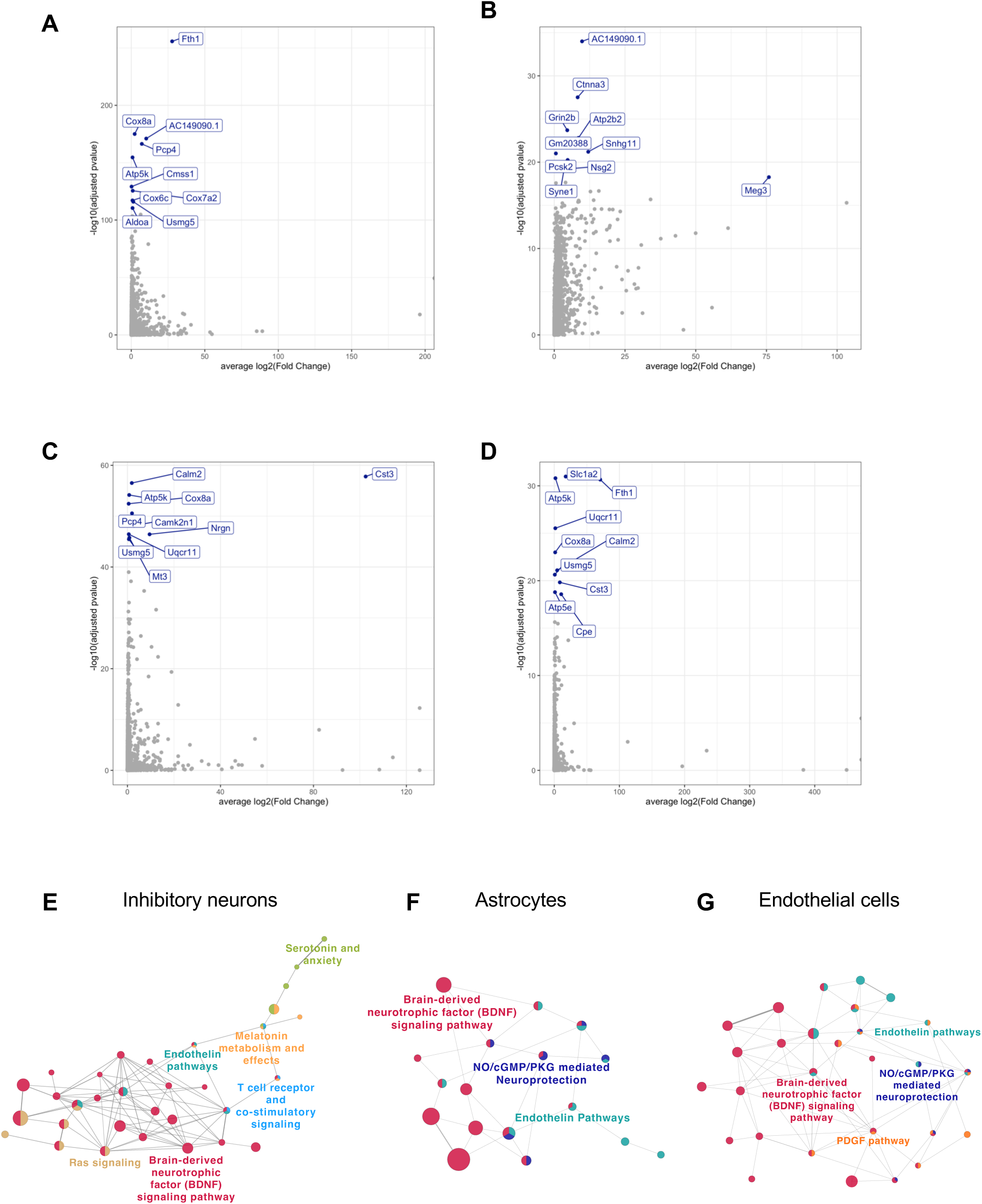
snRNA-seq reveals neuroprotective-induced pathways in forebrain cells. Top-upregulated genes in inhibitory neurons (**A**), *Sv2c*-interneurons (**B**), astrocytes (**C**) and endothelial cells (**D**). Functionally organized network from ClueGO analysis visualized with Cytoscape in inhibitory neurons (**E**), astrocytes (**F**), and endothelial cells (**G**). Only main pathways are represented. Main terms are represented with color. Dot size represents the number of differentially expressed genes in common between pathways.

We next used Cytoscape for pathway interaction network analysis in the clusters of *Sv2c-*interneurons, inhibitory neurons, astrocytes, and endothelial cells. These four clusters showed upregulation of the interaction of the BDNF pathway genes in many signaling pathways as well as the interaction of the BDNF signaling network with other signaling networks (Fig. 3E-G and Supplementary Figs. 6A, B). Inhibitory neurons, endothelial cells, and astrocytes showed interactions between the BDNF and endothelin pathways (Fig. 3F-H). The endothelin signaling pathway regulates cellular proliferation, growth, blood flow, blood pressure, and inflammation in diverse ways from normal to pathological conditions (Yamauchi-Kohno *et al*., 1999). In the endothelial and astrocyte clusters, the endothelin pathways interacted with the NO-cGMP-PKG signaling pathway that promotes vasodilatation, synaptic plasticity, and fear memory consolidation via activation of ERK/MAPK (Ota *et al*., 2008; Francis *et al*., 2010) (Figs. 3G, H). In inhibitory neurons, most pathways identified above by enrichment analysis belong to, or interact with, the BDNF pathway network, except for the pathways of melatonin metabolism/circadian rhythm and serotonin signaling (Supplementary Fig. 6A). Among the four VEGF-C responding clusters, *Sv2c*-expressing neurons displayed the highest number of interactions between the BDNF pathway and other signaling pathways (Supplementary Fig. 6B).

### VEGF-C promotes neuroprotection and immunomodulatory responses in the brain

To identify the subsets of brain cells that may either secrete or directly respond to VEGF-C, we analyzed the distribution of the transcripts of VEGF-C and VEGF-C receptors among the cell clusters (Supplementary Fig. 7). *Vegfc* expression was detected in smooth muscle cells and upregulated in these cells by AAV-VEGF-C administration, in agreement with the previous observation that subsets of perivascular smooth muscle cells are transduced by AAV9 (Antila *et al*., 2017) (Supplementary Fig. 1D). Smooth muscle cells of the brain perivascular spaces may therefore become an additional source of VEGF-C after intra-CSF delivery of AAV-VEGF-C. Among the cell clusters expressing potential receptors for VEGF-C, we found that endothelial cells expressed *Flt4* (VEGFR3) and *Kdr* (VEGFR2) while smooth muscle cells expressed only *Flt4*. Neuropilin2 (*Nrp2)* was detected in neuronal cells including *Sv2c*-expressing interneurons, somatostatin (*Sst)* interneurons, parvalbumin (*Pvalb*) interneurons and pyramidal neurons, as well as in meningeal stromal cells. Integrinα9 transcripts were expressed in OPCs, microglia and perivascular macrophages. The above-mentioned cell clusters are thus potential direct targets of VEGF-C.

### VEGF-C pretreatment ameliorates ischemic stroke outcomes

We first investigated the direct effects of stroke on MLVs and brain tissues in the first week after tMCAO. Quantification of LYVE1+ MLVs showed that they rapidly expanded at 1 day-post stroke onset (1d-pso) (Supplementary Figs. 8A-C). This response was accompanied by an increase of *Vegfc* transcript expression in the ipsilateral hemisphere (Supplementary Fig. 8D). We next tested whether VEGF-C pretreatment may protect mice against ischemic stroke. AAV-VEGF-C and AAV-CTRL mice received tMCAO surgery 4 weeks after administration. (Figs. 4A). We confirmed that the AAV-VEGF-C administration further promoted a robust increase of the LYVE1^+^ MLV coverage and diameter in VEGF-C pretreated mice compared to stroke AAV-CTRL mice at day 7 (Figs. 4B,C).

**Figure 4.**
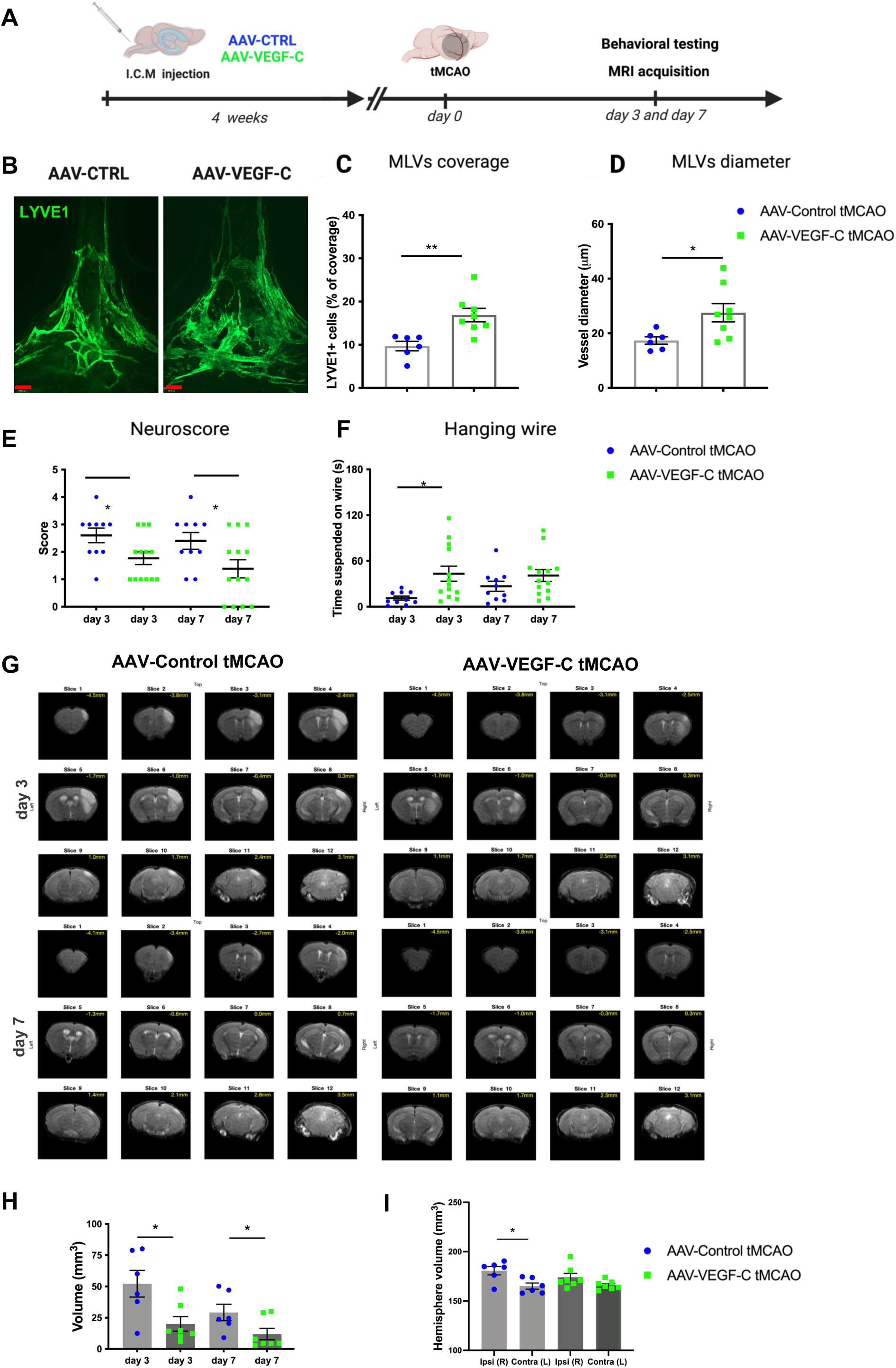
AAV-VEGF-C promotes brain protection, functional recovery and improves adaptive lymphatic growth response to stroke. **(A)** Schematic diagram of the experimental procedure: Mice received an injection of AAV-VEGFC or AAV-CTRL into the cisterna magna and 4 weeks after animals underwent tMCAO. On day 3 and day 7 after tMCAO animals were evaluated using MRI and behavioral tests AAV**-**VEGF-C- increased LYVE1^+^ MLVs growth. **(B)** Anti-LYVE1 immunolabeled MLVs in the confluence of sinuses (COS). VEGF-C-preconditioning increased LYVE1^+^ Quantification of lymphatic coverage **(C)** and diameter (**D**) at 7d-pso (*n* = 6-8 mice/group, \**P* < 0.05, \*\**P* < 0.005); Unpaired t-test, Scale bar: 170 µm.). **(E)** Quantifications illustrating the motor behavior deficit evaluated by the Neuroscore scale, AAV-VEGF-C-pre-treated mice recovered better than controls after tMCAO. **(F)** Hanging wire test evaluation at day 3 post tMCAO. The average time spent on the task and, thereby, the muscular strength was higher in VEGF-C-preconditioned mice than in controls, which reflects less motor impairment (*n =* 10-13 mice/group; \**P <* 0.05; 2-way ANOVA followed by Bonferroni’s multiple comparisons). **(G)** Representative images of MRI anatomical T2 weighted scans showing the infarct lesion induced by tMCAO in AAV-CTRL and AAV-VEGF-C mice. **(H)** Quantification of the lesion volume at days 3 and 7 post tMCAo. AAV-VEGF-C pretreatment promoted a significant reduction of lesion volume when compared to the control treatment (\**P < 0.05;* Wilcoxon test). **(I)** Volumetric quantification of the ipsilateral and contralateral cerebral hemispheres. In control mice, the volume of ipsilateral (Ipsi), or right (R), hemisphere is significantly increased compared to the contralateral (contra) or left (L) hemisphere, indicating cerebral edema in the ipsilateral hemisphere (\**P* < 0.05; *n* = 6-7 animals/group; Wilcoxon test).

tMCAO mice were then subjected to a series of behavioral tests and examined by T2 map and T2-weighted MRI for brain tissue damage, at 3 and 7 d-pso (Fig. 4D). Neurological outcomes were improved in VEGF-C-pretreated mice compared to controls. On the Neuroscore, AAV-VEGF-C treated mice were improved on day 3 and 7 (Fig. 4E). In addition, at 3d-pso, the average time of AAV-VEGF-C treated mice in the hanging wire test was also longer compared to controls (Fig. 4F), which reflects less motor neuromuscular impairment, better muscular strength, and motor coordination at the subacute stage. AAV-VEGF-C treated mice moreover, showed an improving trend in sensorimotor performances measured by the corner test (Supplementary Fig. 9).

*In vivo* MRI of tMCAO mice was used to localize and measure the volume of hyperintense regions corresponding to infarct lesions. As shown in the representative images in Figs. 4G, the infarct lesions were significantly reduced in the AAV-VEGF-C treated compared to control tMCAO mice at day 3 and day 7, which are quantified in Fig 4H. At day 3, VEGF-C pretreated mice also showed a reduction trend in hemispheric edema, as the right hemisphere (ipsilateral to the MCAO occlusion) was not significantly increased, compared to the left hemisphere (contralateral to the MCAO occlusion), in contrast, to control tMCAO mice (Fig. 4I). In AAV-VEGF-C treated mice, MLV expansion thus correlates with the improved sensory-motor behavior and the reduced tissue damage observed after stroke.

### VEGF-C pretreatment protects brain tissues from ischemic insult

After MRI and behavioral studies, brain tissues were isolated from CTRL and VEGF-C pretreated stroke mice and analyzed for histological and gene expression changes induced by VEGF-C. Immunolabeling was performed on 7d-pso brain sections to detect the vascular cells and astrocytes that form the blood-brain barrier, the microglial cells that regulate tissue inflammation, and the neurons. Similar coverage of podocalyxin^+^ blood vessels (PDLX^+^ area) and astrogliosis (GFAP^+^ area) were found in AAV-VEGF-C and AAV-CTRL tMCAO mice (Figs. 5A, B). In contrast, the number of Iba1^+^ microglia and macrophages were strongly reduced in VEGF-C pretreated mice compared to controls (Fig. 5C). Although ischemic infarct volume was reduced by VEGF-C pretreatment (Figs. 4D, E), the density of perilesional NeuN^+^ neurons was not significantly increased compared to controls (Fig. 5D). These results suggest that the neuroprotective effect of VEGF-C was not mediated by angiogenesis or enhanced neuronal survival, but perhaps by mitigating inflammatory responses.

**Figure 5.**
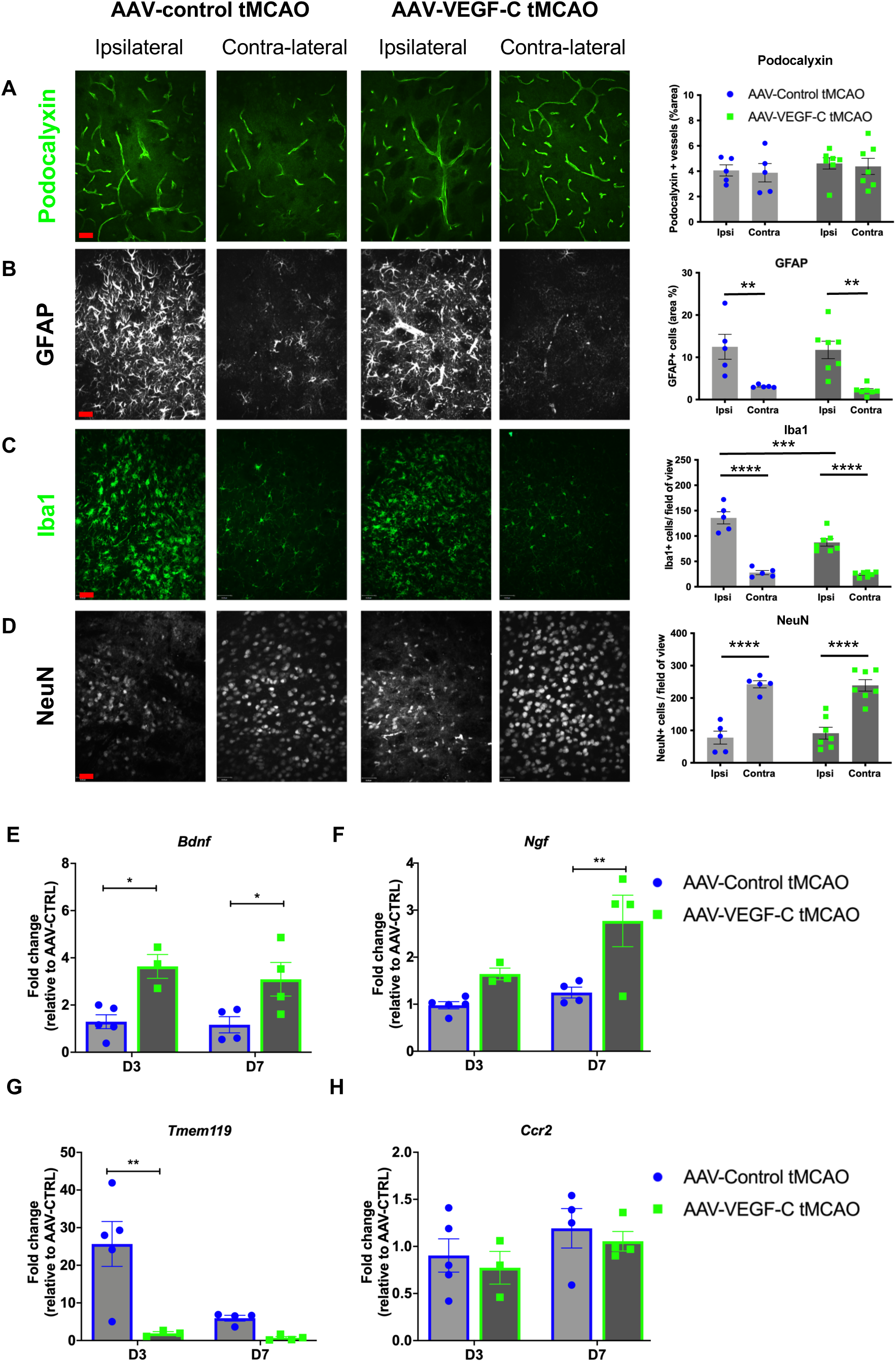
AAV-VEGF-C administration before ischemic stroke increases the expression of neurotrophins and prevents microglia expansion at 7d-pso. **A-D** Representative images and quantifications of brain immunostaining in the ipsilateral (Ipsi) and contralateral (contra) hemispheres of AAV-Control and AAV-VEGF- C injected mice 7 days after tMCAO. **(A)** AAV-VEGF-C preconditioning did not induce cerebral angiogenesis in the lesion border on day 7 (%) for PDLX^+^ blood vessels. **(B)** Astrogliosis (GFAP^+^ astrocytes) was induced in the ipsilateral hemisphere, as shown in representative images taken in the lesion border in the striatum, but with a similar extent in control and VEGF-C pretreated mice **(C)** Quantification of microglia/macrophages expressing ionized calcium-binding adaptor molecule 1 (Iba1+). The number of Iba1+ cells/ field of view was significantly reduced in AAV-VEGF-C mice compared to controls. **(D)** NeuN^+^ neurons were significantly reduced in number in the ipsilateral compared to the contralateral hemisphere due to the tMCAO injury (\*\*\*\**P* < 0.0001), but no difference was observed between AAV-VEGF-C mice and controls. One-way ANOVA and Bonferroni’s post hoc test. Data are represented as mean ± S.E.M, *n* = 5-7 mice in each experimental group. q-PCR analysis of forebrain homogenates (ipsilateral hemisphere) at days 3 and 7 after stroke shows that mRNA expression of **(E)** Brain-derived neurotrophic factor (*Bdnf*) and **(F)** Nerve growth factor (*Ngf*) genes were increased in the group tMCAO AAV-VEGF-C compared to controls. The expression of **(G)** transmembrane protein 119 (*Tmem119*) gene, a microglia specific-cell surface marker, was highly downregulated at day 3 post-stroke in the AAV-VEGEF-C group as compared to controls, **(H)** while expression of macrophages/monocytes marker (*Ccr2*) was similar between the two groups. Scale bar: 35µm

Using Q-PCR, we next investigated the molecular signals that may account for microglial and neural cell behavior at the early and later stages after stroke (3d- and 7d- pso). VEGF-C pretreatment was associated with a persistent increase in the transcription of neuroprotective genes such as *Bdnf* and *Ngf* (Figs. 5E, F). The gene expression signatures of microglia (*Tmem119*) and macrophages/monocytes (*Ccr2*) were analyzed to better understand the respective contribution of resident microglia and circulating myeloid cells to the Iba1^+^ cell population quantified on brain sections. VEGF-C pretreatment resulted in a strong reduction of *Tmem119* expression (FC = 10 at 3d-pso) compared to control mice, without change in *Ccr2* expression (Figs. 5G, H). Thus, the reduction of Iba1^+^ cells observed in VEGF-C pretreated mice thus mainly suggests reduction of microglia rather than macrophages. Interestingly, VEGF-C pretreatment increased the expression of MHC-II (Supplementary Fig. 10A), and thereby suggesting activation of microglia and adaptive immune responses. Moreover, we found increased expression of the T lymphocyte marker (*Cd3*) and the chemokine receptor type 7 (*Ccr7*) which regulates T cell migration (FC = 4 for *Cd3*, and 2.5 for *Ccr7* at 3d-pso) (Supplementary Figs. 10B, C), suggesting increased infiltration of circulating lymphocytes into the injured brain tissues.

The transcription of genes encoding pro-inflammatory cytokines (*Il6*, *Il12A*, *Tnfa)* and the secreted pro-apoptotic molecule granzyme B (*Grzb*) was increased at day 3 after VEGF-C treated mice (Supplementary Figs. 10D-G). Expression of Interferon-gamma (*Ifng*), a major pro-inflammatory cytokine gene induced by stroke, was strongly repressed (FC = 4.5) at 7d-pso in AAV-VEGFC-treated mice (Supplementary Figs. 10H). The se results demonstrate that the kinetics of pro-inflammatory molecule expression after stroke is changed upon in AAV-VEGF-C treatment. In contrast, the expression of anti-inflammatory cytokine genes (*Il-4*, *Il-10*, *Tgfb1*) and genes related to anti-inflammatory polarization of microglia/macrophages (*Mmp9*, *Mrc1*) was similar in VEGF-C pre-treated and control mice (Supplementary Figs. 10i-m). We furthermore found that VEGF-C pre-treatment reinforced the ligand/receptor signaling between CCL3 and ACKR2 (Supplementary Figs. 10N, 10O). Therefore, the above postmortem data revealed multiple effects of VEGF-C pretreatment, including neuroprotection and a complex regulation of neuroinflammation in forebrain tissues after stroke. Altogether these VEGF-C-induced responses may protect against tissue damage and correlate with an improvement in outcomes after ischemic stroke injury.

Finally, we tested if the protective effects of VEGF-C could be used as a therapy. We administrated recombinant protein rVEGF-C intracisternally following tMCAo immediately after reperfusion. The animals were evaluated using the same protocol as the group treated pre-stroke, including behavioral testing, MRI evaluation, and histological analysis of meningeal lymphatics. Interestingly, VEGF-C treatment at reperfusion had no effect on mLV coverage (Fig. 6A-C). We also observed that rVEGF-C treatment at this time-point had no protective effect regarding the functional outcome or lesion size (Fig. 6D-I), suggesting that the effects observed upon AAV-VEGF-C pretreatment are largely dependent of the expansion of mLVs and the long-term effects of VEGF-C signaling.

**Figure 6.**
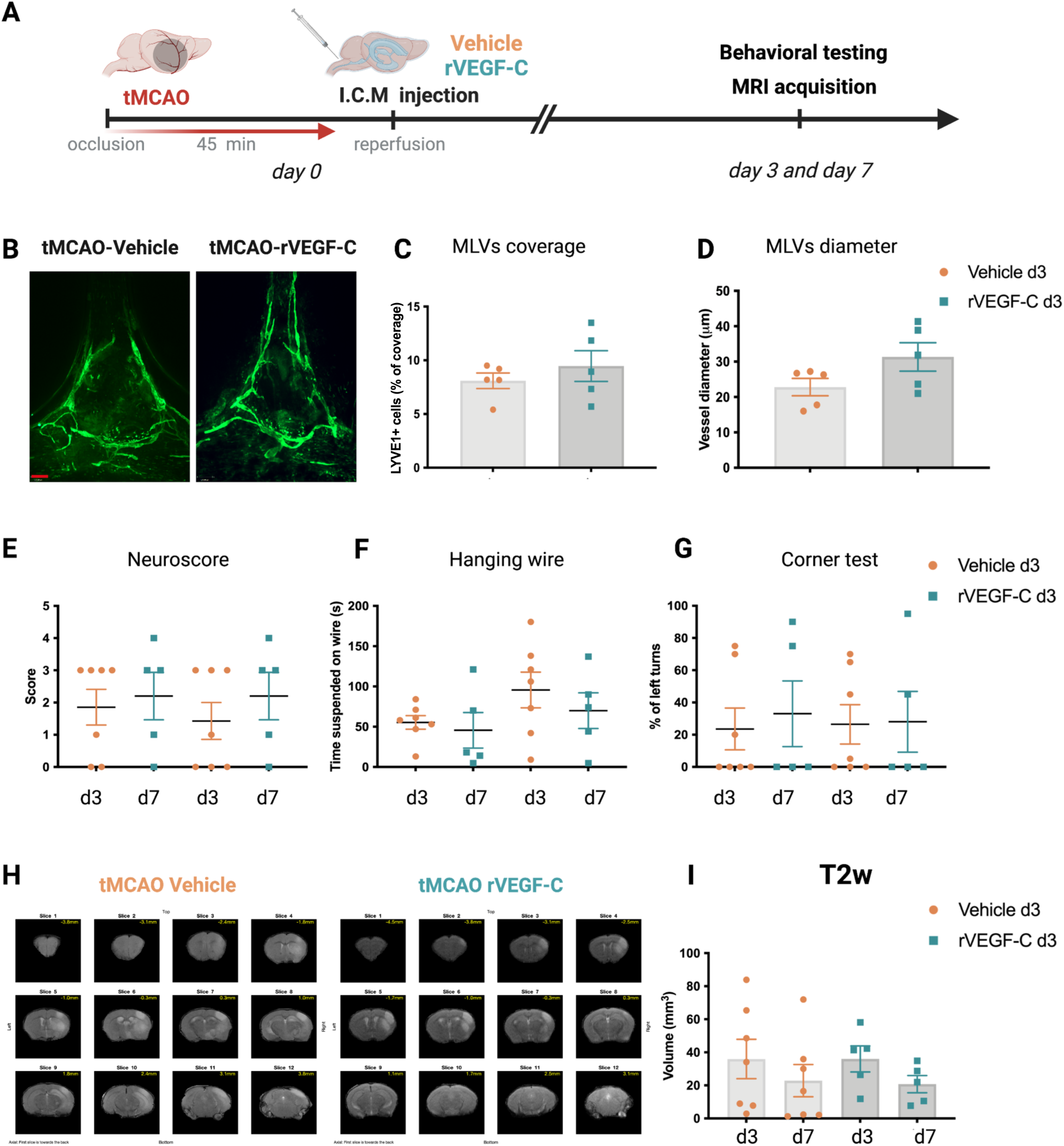
Single-dose treatment with rVEGF-C does not improve outcomes after tMCAO. **(A) Experimental setting:** Mice underwent tMCAO and, after reperfusion, received an ICM injection of either recombinant VEGF-C (Cys156Ser protein, rVEGF-C) or vehicle control (0,25% BSA). **(B)** Anti-LYVE1 immunolabeled MLVs in the confluence of sinuses (COS) at day 7 post tMCAO. Quantification of MLV surface shows that rVEGF-C did not affect MLV coverage **(C)** or MLV diameter **(D)** (*n* = 5 mice/group, Unpaired t-test). **(E)** Neuroscore scale shows a similar score between rVEGF-C or vehicle group. **(F)** A similar time of suspension in the hanging wire test. (**G)** No difference in the percentage of left turns (impaired side) in the Corner test at day 3 and day 7 (*n* = 7, 5 mice/group one-way ANOVA test). **(H)** Representative image of MRI anatomical T2 weighted scans showing the infarct lesion. **(I)** Quantification of the lesion volume at days 3 and 7 post stroke (*n* = 7-5 mice/group Kruskal-Wallis test) rVEGF-C treatment did not influence the lesion volume when compared to vehicle. Scale bar: 170 µm .

## Discussion

Here, we have used cutting-edge techniques to investigate the impact of stimulating VEGF-C signaling using intra-cisternal administration of AAV-VEGF-C on CNS/ISF solute drainage and brain cell transcriptional programs both in physiological conditions and after acute ischemic stroke. First, the combination of intra-CSF administration of Gd-DOTA with anatomical MRI and quantitative T1 mapping in live mice demonstrated that AAV-VEGF-C promotes lymphatic drainage to dCLNs without affecting the glymphatic pathway. This real-time observation is in agreement with reports on post-mortem studies showing that the ICM injection of either AAV-VEGF-C or VEGF-C promotes the expansion of mLVs and increases the transport of CSF fluorescent molecules, nanoparticles, or brain tumor antigens from the CSF into the brain-draining cervical LNs (Da Mesquita *et al*., 2018; Song *et al*., 2020). Second, snRNA-seq data demonstrated transcriptomic changes in several brain cell clusters stimulated by AAV-VEGF-C, including support of synaptic function, BDNF, and calcium signaling in a neuroprotective signature. AAV-VEGF-C administration into the CSF has thus dual effects on brain cells and brain-draining LNs, as a logical consequence of AAV-VEGF-C transport via both the glymphatic system and the lymphatic CSF/ISF outflow pathway.

Intracisternal VEGF-C administration is well known to expand and enlarge lymphatic vessels in the meninges (Aspelund *et al*., 2015; Louveau *et al*., 2015; Antila *et al*., 2017; Song *et al*., 2020) and the cribriform plate tissues (Hsu *et al*., 2020). In the extracerebral spaces, we found that AAV-VEGF-C also stimulated the growth of extracranial lymphatics that collect into mandibular or cervical LNs, without inducing lymphangiogenesis in distant head tissues such as the ear. The lymphangiogenic effect of AAV-VEGF-C thus propagates along the CNS draining lymphatic vasculature toward brain-draining LNs (Esposito *et al*., 2019). We were unable to obtain direct MRI evidence that CSF/ISF outflow through the cribriform plate was augmented by AAV-VEGF-C treatment, suggesting that additional lymphatic circuits, for example in the basal skull, were stimulated by VEGF-C and synergized with cribriform lymphatics for the benefit of increased dCLN drainage. We also noted that glymphatic transport as evidenced by a one-time ‘snapshot’ T1 map ∼1hr after CSF contrast administration was not altered in AAV-VEGF-C treated mice. In the present experimental setting, solute transport via the glymphatic system was thus not promoted by VEGF-C, either directly or indirectly via its downstream effect on lymphatic vasculature growth.

We have characterized brain transcriptomic responses to the intracisternal delivery of AAV-VEGF-C from a large sample of brain nuclei (78,728) that were analyzed by shallow sequencing (a median of 1,627 genes and 2,550 transcripts per nucleus) and further comprehensive integration of single-nucleus data (Stuart *et al*., 2019). We focused on the most prominent and statistically significant changes induced by AAV-VEGF-C treatment. This approach however precluded the identification of small-size subpopulations of brain cells such as neural stem/progenitor cells and immune cell subtypes and characterization of downstream effectors in altered signaling pathways. Despite this limitation, we found that intracisternal AAV-VEGF-C administration impacted the function of different brain cell subtypes as well as the expression of *Vegfc* and VEGF-C receptor genes. The most significant intracellular signaling alterations caused by AAV-VEGF-C treatment were observed in inhibitory neurons and *Sv2c*- interneurons. GO annotations showed that AAV-VEGF-C induced an increase in calcium signaling in these cell clusters. Genes encoding calcium signaling regulators such as *Pcp4, ATP2B2, GRIN2B,* and *Calm2* also among the top-upregulated genes by VEGF-C in interneurons. In neural cells, Ca^2+^ is an essential messenger molecule that regulates synaptic activity through cell depolarization, vesicle fusion, and vesicle recycling (Clapham, 2007). Thus VEGF-C may thus positively impact inhibitory interneuron activity and synaptic plasticity.

In the context of stroke, we investigated the lymphatic response in parallel with the brain damage and neurological outcomes after a transient ischemic injury. We confirmed that ischemic stroke induced an acute increase in MLV coverage and caliber but only transiently within 24 hours. CSF participates in the acute swelling of ischemic brain tissues, by entering within minutes of the stroke insult along perivascular flow channels (Mestre *et al*., 2020). This massive increase in fluid within the brain leads to unbalanced CSF circulation, necessitating readjustment of CSF drainage which may explain the acute MLV increase shortly after stroke. MLV outgrowth after stroke injury also correlated with a surge of cortical *Vegfc* expression, which confirms previous reports of upregulated VEGF-C protein expression in brain cells after ischemia, either within 24h after focal photo-thrombosis (Jiang and Liao, 2010) or 72h after a 5 minutes ischemia (Bhuiyan *et al*., 2015). VEGF-C produced by brain tissues surrounding the lesion may thus contribute to MLV outgrowth via the glymphatic transport of ISF/CSF solutes.

The effects of VEGF-C on stroke outcomes were first evaluated after pretreatment by ICM injection of AAV-VEGF-C. VEGF-C improved the functional recovery and reduced the lesion size evaluated with MRI, at 3 and 7d-pso, as compared to AAV-CTRL treatment. This protective effect of AAV-VEGF-C pretreatment may have several causes. One explanation is provided by snRNA-seq data analysis showing that VEGF-C triggers neurosupportive pathways, especially calcium and BDNF signaling, in interneurons which may facilitate resistance to ischemic injury. VEGF-C also induced meningeal and extracranial lymphatic growth as well as increased dCLN drainage which is expected to improve the clearance of brain lesion-derived fluids and debris. In support of this explanation, K14-Vegfr3-Ig transgenic mice with reduced lymphatic vasculature displayed larger brain lesions than controls after ischemic stroke (Yanev *et al*., 2020). VEGF-C-induced lymphatic growth may moreover promote immune cell trafficking and modulation of the inflammatory response in the injured brain tissues. VEGF-C has been demonstrated to have anti-inflammatory effects in a variety of disease models, including inflammatory bowel disease (IBD), rheumatoid arthritis, and skin inflammation (Schwager and Detmar, 2019). Our present results are in line with this set of studies and suggest anti-inflammatory potential of VEGF-C in our experimental stroke model. VEGF-C driven lymphatic growth can however increase brain antigens’ drainage to dCLNs and thereby contribute to peripheral immune activation (Song *et al*., 2020). Along this line, brain-to-CLN signaling has been reported to trigger systemic inflammatory responses after acute stroke (Esposito *et al*., 2019). In this study, administration of MAZ51, a tyrosine kinase inhibitor of VEGFR-3 and VEGFR-2, or surgical removal of superficial cervical LNs, led to significant reduction of infarct at 72h after cerebral ischemia, although no statistically significant differences in neurological recovery were detected. These loss-of VEGF-C signaling experiments however differ from the direct increase of VEGF-C induced by AAV-VEGF-C administration. They also not reflect on the neurosupportive effect of AAV-VEGF-C on brain cells which may explain that our experimental model showed no deleterious effect of VEGF-C and improved neurological recovery.

The combination of histological and gene expression analysis of brain tissues after stroke reveals some mechanisms that may be activated by VEGF-C. AAV-VEGF-C pretreatment resulted in a reduced population of brain Iba1^+^ cells at 7d-pso, which correlated with the reduction of brain *Tmem119* transcripts in mice with ischemic stroke. The expression of *Ccr2*, a marker of circulating monocytes/macrophages, was however not altered in these mice. In ischemic stroke, activated microglia play a pivotal role in the initial immune response (Ma *et al*., 2017), producing pro-inflammatory cytokines such as TNF-alpha and nitric oxide that ultimately contribute to the release of free radicals and mitochondrial damage (Lambertsen *et al*., 2009). VEGF-C thus repressed the expansion of microglial population without increasing infiltration of circulating macrophages, which may provide immunoprotection against post-ischemic inflammation. Interestingly, this protective effect of VEGF-C may be indirectly mediated by MLVs, as impairment of meningeal lymphatic drainage can exacerbate microglial inflammatory response, as suggested in a model of Alzheimer’s disease (Da Mesquita *et al*., 2021).

A mild activation of microglia could however favor neuroplasticity via the secretion of neurotrophic factors such as BDNF and NGF (Elkabes, DiCicco-Bloom and Black, 1996). BDNF is the most abundant neurotrophin expressed in the adult brain and has survival- and growth-promoting actions on a variety of neurons (Wang *et al*., 2014), while NGF regulates neuronal plasticity (Rocco *et al*., 2018) and neuroinflammatory responses (Minnone, De Benedetti and Bracci-Laudiero, 2017). Interestingly, AAV-VEGF-C pretreatment also upregulated the expression of *Bdnf* and *Ngf* transcripts after stroke injury, which may support neural cell activity and add to the immunomodulation meditated by VEGF-C during post-ischemic inflammation.

Finally, we tested if VEGF-C could induce similar protective effects when administrated after tMCAO. Unfortunately, a single dose of recombinant protein administration failed to show the same beneficial effects observed after the AAV-VEGF-C administration before the ischemic insult. A possible reason for this outcome is the short half-life of VEGF recombinant proteins (Rennel *et al*., 2008) and the relatively low amount of injected VEGF-C which even failed to induce MLV outgrowth (Supplementary Fig. 8). One study using a biomaterial (hydrogel hyaluronic acid or methylcellulose-based) for rVEGF-C delivery has reported a sustained release of VEGF-C up to 7-10 days *in vitro* (Louveau *et al*., 2015). Such approach could be tested in the future to determine the efficacy of VEGF-C as a potential post-stroke therapy.

The present study brings new insights into the range of effects induced by intra-CSF delivery of AAV-VEGF-C, including lymphatic growth along the CSF/ISF outflow pathway, increased drainage of CNS-derived fluids in dCLNs, neuroprotective responses in neural cells and immunomodulation of microglial cells.

## Methods

### Study approval and mice

All *in vivo* procedures used in this study complied with USA federal guidelines and the institutional policies of the Yale School of Medicine Animal Care and Use Committee. Male C57BL/6J mice (purchased from Jackson Laboratory) aged 4 to 10 weeks old were used in the present study.

### Intracisterna-magna (ICM) injections

For ICM injections, mice were anesthetized by intraperitoneal (i.p) injection of ketamine (80 mg/kg), and xylazine mix (20 mg/kg) the dorsal neck was shaved and cleaned using povidone-Iodine scrub. After positioning the mouse in the stereotactic apparatus, a 1 cm incision was made at the base of the skull, and the dorsal neck muscles were separated using forceps. AAVs serotype 9 (AAV_9_) were administered by ICM injection. A single dose of 2μl (3 x 10^9^ viral particles) of AAV_9_-YFP, AAV_9_-mouse VEGF-C (AAV-VEGF-C) or AAV_9_-mouseVEGFR34–7-Ig (AAV-CTRL) was administered into young (4 weeks old) C57BL/6J male mice. For the post-stroke experiment, a single dose of 1μg of recombinant protein VEGF-C^156S^ was diluted in 2μl of sterile PBS containing 0.1% BSA (vehicle). ICM injections were performed using a Hamilton syringe with a 34-G needle at a 15 degrees angle. The needle tip was retracted 2 min after the injection. All AAVs were produced by the vectorology platform of Paris Brain Institute (ICM).

### Nuclei Isolation and Library Preparation

To generate transcriptome profiles for mouse brains after AAV-VEGF-C administration we used a total of 10 brain samples from adult male mice (8 weeks old). Five brains from AAV-VEGF-C and AAV-CTRL mice were collected, cryopreserved, pooled, and then processed using the 10x Genomics protocol.

Microfluidic capturing of nuclei and libraries were prepared according to manufacturers’ protocol (10x Genomics; Chromium Single Cell 3L GEM, Library & Gel Bead Kit v3, 16 rxns PN-100007)

### Sequencing of libraries

The sequencing was performed by following manufacturers direction (10x Genomics; CG000183) for targeted 30000 reads per nucleus, on HiSeq 4000 platform (Illumina). In total, we sequenced 78,728 nuclei profiles from snRNA-seq, with a median of 1,627 genes and 2,550 transcripts per nucleus (Supplemental Table 1). MEX files obtained were then loaded into Seurat, cells presenting less than 200UMI were discarded, and the samples were normalized using the SCTransform function with regression of the mitochondrial RNA percentage. Batch effects were corrected using the Seurat integration pipeline (Butler *et al*., 2018) on the 5,000 most variable genes. We then aggregated transcriptionally similar cells, using the Seurat t-distributed stochastic neighbor embeddings (tSNE) algorithm. Next, we removed clusters likely to be of low quality, resulting from debris, doublets/multiplets and dead cells by regressing out the clusters expressing less than 1,000 counts on average to further enhance the quality of our analysis.

The 78,728 nuclei remaining after processing were clustered using the FindCluster function with k-nearest neighbors of 15, and a resolution of 1.2. The clusters were then manually identified according to their top markers with the help of the Allen Brain atlas (*Reference Atlas :: Allen Brain Atlas: Mouse Brain*, no date). A sub-clustering of the identified endothelial, microglial/macrophage clusters was then realized using a resolution of 0.8 of the SNN matrix object, with the same default k-nearest neighbors (Waltman and Eck, 2013).

Differential expression analysis was conducted using FindMarkers function from the Seurat pipeline, using a Wilcoxon test for p-value estimation with a Bonferroni correction for the number of tests. Only differentially up-regulated genes (adjusted p-value above 1.5) were used for enrichment analysis. The enrichment analysis was performed with EnrichR R packages on Wikipathways database. We selected pathways with at least 20 genes and with a p-value under 0.05. Cytoscape (ClueGO®) application was used to visualize enrichment analysis results into functionally organized networks.

### Contrast agent-labeled CSF MRI for glymphatic-lymphatic transport

All the MRI experiments were conducted at Magnetic Resonance Research Center at Yale University. T1 mapping was used to evaluate glymphatic transport and brain-derived fluid drainage to the cervical lymph nodes after VEGF-C administration. MRI acquisitions were performed on a Bruker 9.4 T/16 MRI scanner with a BGA-9S-HP imaging gradient interfaced to a Bruker Advance III console and controlled by Paravision 6.1 software (Bruker Bio Spin, Billerica, MA, USA). The 3D T1 mapping technique used in this study followed the previously published protocol (Xue *et al*., 2020).

For the CSF contrast administration and MRI procedures all mice (*n* = 20, 10 injected with AAV-VEGF-C and 10 with AAV-CTRL), were anesthetized with a ketamine/xylazine (KX) mixture: (ketamine 17.5 mg/ml and xylazine 2.5 mg/ml, 0.1ml/20g body weight) combined with glycopyrrolate (0.2mg/kg IP). Anesthesia was maintained with KX (0.05 ml of the KX-mixture/20g body weight) administered every 30 minutes via an IP catheter and supplemented with a 1:1 air/O_2_ mixture. CSF administration of gadoteric acid (Gd-DOTA, Guerbet LLC, Princeton, NJ, US) was performed via the cisterna magna as previously described (Xue *et al*., 2020). Briefly, we used a 34-ga needle connected via polyurethane tubing to a 50µl Hamilton syringe (Hamilton, US) mounted in a micro-infusion pump (Legato 130, KD Scientific, Holliston, MA, USA), MW 559 Da). Seven µl of Gd-DOTA prepared as a 1:20 dilution in sterile 0.9% NaCl was administered at an infusion rate of 1 µl/min. After the CSF infusion, the anesthetized mouse was transferred to the 9.4T MRI for T1 mapping. The T1 scan was pre-fixed to start 50min from initiation of CSF infusion on the bench. For T1 maps covering the head and neck of the mouse was acquired using two steps: First, a spatial inhomogeneity profile of the RF transmit (B1+) was acquired using a double angle method using a rapid acquisition with relaxation enhancement (RARE) sequence (TRL = 10000 ms, TEL =L 22 ms, AverageL =L 1, RARE factorL = 4, number of slicesL = 36, in plane resolutionL = 0.24 mm/pixel, slice-thicknessL = 0.3 mm, slice gapL = 0.2 mm Flip angles = 70° and 140°). Second, a spoiled gradient echo VFA-SPGR method (TRL=L16 ms, TEL=L3 ms, AverageL=L1, scanning timeL=L2 min 40 s, matrixL=L100L×L100x100 reconstructed at 0.18L×L0.18x0.18 mm). A set of six flip angles (2°, 5°, 10°, 15°, 20°, 30°) was used and the total scan requiredL∼L16 min. The T1 maps were filtered and T1 values > 5000ms were excluded. MRIs comprising the summed low flip angle (2° and 5°) SPGR images were used as anatomical templates for outlining the brain and lymph nodes. The T_1_ maps of the head were used to quantify the volume of glymphatic transport which was defined as brain tissue voxels with a T1 in the range of 1-1700ms (this particular T1 range represents tissue which has been shortened by the uptake of Gd-DOTA). For each mouse, the glymphatic transport volume was extracted using PMOD software (PMOD, version 4.0). Similarly, outflow to the nasal conchae and drainage to the deep cervical lymph nodes were quantified by first outlining these anatomical structures and then extracting voxels with T1 values from 1-1700ms.

### Focal cerebral Ischemia

Focal cerebral ischemia–reperfusion was induced as described previously (Longa et. al 1989). In brief, transient focal ischemia was induced under ketamine (80 mg/kg IP) and xylazine (10 mg/kg IP) anesthesia. Body temperature was maintained at 37.0±0.5°C throughout the procedure with help of a homeothermic monitoring system (Harvard Apparatus 55-7020, Massachusetts, USA). After a midline neck incision, the right external carotid and pterygopalatine arteries were isolated and cauterized. The internal carotid artery was lifted and occluded at the peripheral site of its bifurcation as soon as the distal common carotid artery was clamped. Focal cerebral ischemia was induced by intraluminal filament occlusion of the right middle cerebral artery (MCA) for 45 minutes using a 6-0 nylon monofilament with a silicone-coated tip (Doccol Co., Redlands, CA). Reduction in regional cerebral blood flow was confirmed using trans-cranial laser-Doppler flowmetry (MoorVMS-LDF1, Moor instruments, Delaware, USA) in the cerebral cortex supplied by the MCA. Sham-operated mice were anesthetized, and the common carotid artery was dissected free from surrounding connective tissue, but the MCA was not occluded.

### Behavioral testing

Three behavioral tests were used to functionally assess the sensorimotor function after tMCAO surgery. The behavioral tests were conducted by an evaluator blinded to the experimental groups. The behavioral tests were performed on day 3 and day 7 post-tMCAo surgery. Neurological deficit was evaluated using a 5-point scoring system (0, no deficit; 1, forelimb weakness and torso turning to the ipsilateral side when held by the tail; 2, circling to one side; 3, unable to bear weight on affected side; and 4, no spontaneous locomotor activity or barrel rolling) as described previously.

The hanging wire test evaluates both limb strength and balance after MCAO. The apparatus consists of a 50 cm wide 2-mm thick metallic wire, secured to two vertical stands at around 30cm above a foam pillow. The time-out period was 180 seconds for each trial. The average time of suspension in three different trials with 5 min rest was then calculated.

The corner is commonly used for identifying and quantifying sensorimotor and postural asymmetries (Zhang *et al*., 2002). The apparatus consists of two acrylic boards placed closely together at a 30-degree angle forming a narrow alley. A mouse is then placed in between the boards facing the corner. As the animal approaches the corner, both sides of the vibrissae are simultaneously stimulated which leads the animal to rear and turn 180 degrees. Animals with unilateral brain damage will preferentially turn around in the ipsilateral direction (non-impaired side). The percentage of left turns in a total of twenty trials was calculated.

### Measurement of infarct volume

MRI-based translational imaging was used to determine the infarct volume after tMCAO. A cohort of animals was evaluated at 3d-pso and again at 7d-pso. Briefly, MRI data were obtained on a modified 11.7T system with a Bruker spectrometer (Bruker Bio Spin, Billerica, MA, USA). Mice were anesthetized with 2% isoflurane and maintained in a mixture of 1% isoflurane, 30% of O2, and 70% N2O using a nose cone. Respiration rate (50–80 breaths/min), and rectal temperature (37 ± 1°C) were continuously monitored and maintained using warm water-pumped system. MR Images were acquired using a Transmit-only volume (70 mm birdcage) coil and receive-only surface (35 mm diameter ring) coil configuration (RAPID MR international)). First, anatomical images were acquired using a T2-weighted RARE sequence with 2 averages and a RARE factor of 4 (TE = 24 ms, TR= 3000 ms, FOV: 32 x 32, Matrix: 128 x 128, slice thickness: 1mm, number of slices = 9). T2 measurements were acquired using a Multi-Slice Multi-Echo (MSME) sequence (number of echoes = 10, TR:LL3000 ms, TE (ms): 10, 20, 30, 40, 50, 60, 80, 90 and 100, matrix: 85x85, FOV: 17x17, slice thickness: 0.7mm and number of slices: 12. A custom written script using MATLAB version R2019b (MathWorks) was used to fit the data and generate R2 (i.e., 1/T2) maps from the acquired data. The infarct lesion was defined in a semi-automatic fashion. The anatomical images were used to exclude ventricles and to delimitate the brain tissue using MATLAB (Natick, MA). Following we use BioImage suite software (yale.edu/bioimaging/suite) to detect hyperintense regions on the T2-map using a threshold. The total stroke volume was calculated as the sum of the voxels included in the hyperintense regions across all slices, multiplied by the total slice thickness.

### Tissue preparation

Mice were deeply anesthetized with isoflurane and then perfused transcardially with cold phosphate-buffered saline (PBS), followed by 4% paraformaldehyde (PFA). Head dissection was done as described previously (Antila *et al*., 2017). The dissected skullcaps with the dura mater were post-fixed overnight, washed in PBS, and processed for staining.

### Immunolabeling of whole-mount preparations and brain sections

For whole-mount staining of the meninges, the fixed tissues were blocked with 10% donkey serum 2% bovine serum albumin (BSA), 0.5% PBS-Triton-X (blocking solution) overnight. Primary antibodies were diluted in DIM, and samples were incubated in the primary antibody mix at least overnight at 4°C. After washes with PBS-Triton-X in room temperature, the tissues were incubated with fluorophore-conjugated secondary antibodies in PBS-TX overnight at 4°C, followed by washing in 0.5% PBS-Triton-X at RT. After post-fixation in 1% PFA for 5 min, and washing with PBS, the stained samples were transferred to PBS containing 0.02% sodium azide at 4°C and imaged as soon as possible.

For floating sections of the brain, the brain samples were removed and post-fixed overnight. Sections of 30um thick were obtained using a vibratome. Sections were incubated in a blocking solution for 2 h at room temperature, followed by overnight incubation with primary antibodies. After washing with 0.3% PBS-TX three times, the sections were incubated with fluorescent dye–conjugated secondary antibodies in PBS with 3% donkey serum for 2 h at room temperature. After three washes with 0.3% PBS-Triton-X, the sections were mounted with fluorescent mounting medium (Dako) between a glass slide and cover glasses.

For cryosections of LNs (LNs), the LNs were collected, post-fixed overnight in 4% PFA and cryopreserved in 30% sucrose. Sections (20μm) were obtained using a cryostat (model). For cryosections of the head to detection of olfactory bulb surrounding lymphatic vessels, the fixed tissues underwent decalcification with 0.5 M EDTA, pH 7.4, at 4°C for 7 days as described previously (Antila *et al*., 2017) Samples were washed with PBS and immersed in PBS containing 20% sucrose for 24h at 4°C, embedded in OCT compound (Tissue-Tek), and frozen for storage at −80°C. Sections of 30um thick were obtained and the samples were immediately mounted in a glass slide. Following the sections were immunostained using the same protocol as described above.

### Antibodies

The complete list of primary and secondary antibodies and concentrations is in the supplementary material.

### Confocal microscopy: image acquisition and analysis

Laser scanning confocal images of the fluorescent labeled brain and whole-mount skullcap mice were acquired using a spinning-disk confocal (Nikon Eclipse Ti) microscope or a SP8 microscope.

Quantitative analysis of meningeal lymphatic coverage and diameter was performed using FIJI Image-processing software (National Institutes of Health). The percentage of the area covered by LYVE1+ MLVs was detected and quantified semi-automatically in the confluence of sinuses. The diameter of MLVs was calculated by the total area of the vessel/length, as described previously (Zhang *et al*., 2018). The mean value of 8 different segments of the MLVs was then calculated.

For the brain samples, the number of cells was manually counted for microglial cells (Iba1+), and neurons (NeuN+). The percentage of area occupied by GFAP+ or PDLX+ cells was calculated in a semi-automatic fashion using FIJI.

### Statistical analyses

All graphs and statistical analyses were produced using GraphPad Prism 8. Results were expressed as meanL±Lstandard error of the mean. Experiments were performed with full blinding, allocation concealment, and randomization. The data normal distribution was evaluated using the Shapiro-Wilk test. When only two groups were compared, an unpaired t-test was used, the Mann-Whitney test or Wilcoxon analysis. Multiple two-way ANOVA. *P*J<L0.05 was statistically significant.

## Acknowledgments

This work was Supported by grants from the National Institute of Health (NIBIB R01EB016629-01 to JLT, LB, SL; P30 NS-052519, R01 MH-067528 and R01 EB-023366 to FH), Agence Nationale de la Recherche (ANR-17-CE14-0005 BrainWash to JLT, AE), ERC advance (AE, LG). We thank Nadège Sarrazin from the PHENO-ICMice that is supported by ANR-10-IAIHU-06, ANR-11-INBS-0011-NeurATRIS and the Fondation pour la Recherche Médicale.

## Authorship

Study concept: JLT, LB, SL

Study coordination and manuscript writing: JLT, LB, JB, LG, AE Figures: LB, JB, MP

tMCAO surgery: SL, LB, SZ

RNA-seq: preparation (MS), dataset analysis (JB, JG, ML) DCE-MRI: HB, YX, XL

MRI: MP, BS

WB and qPCR: LG, LB

Provision of reagents: JLT, KA

Interpretation of data: JLT, LB, JB, LG, AE, HB, LS, FH, KA, NS, JZ

Curation of data: LB, HB, JG, ML

Statistical analyses: LB, JB, LG

All authors contributed and reviewed the final manuscript

