## Supplementary figures and images for "VEGF-C promotes brain-derived fluid drainage, confers neuroprotection, and improves stroke outcomes"

### Supplementary fig 1

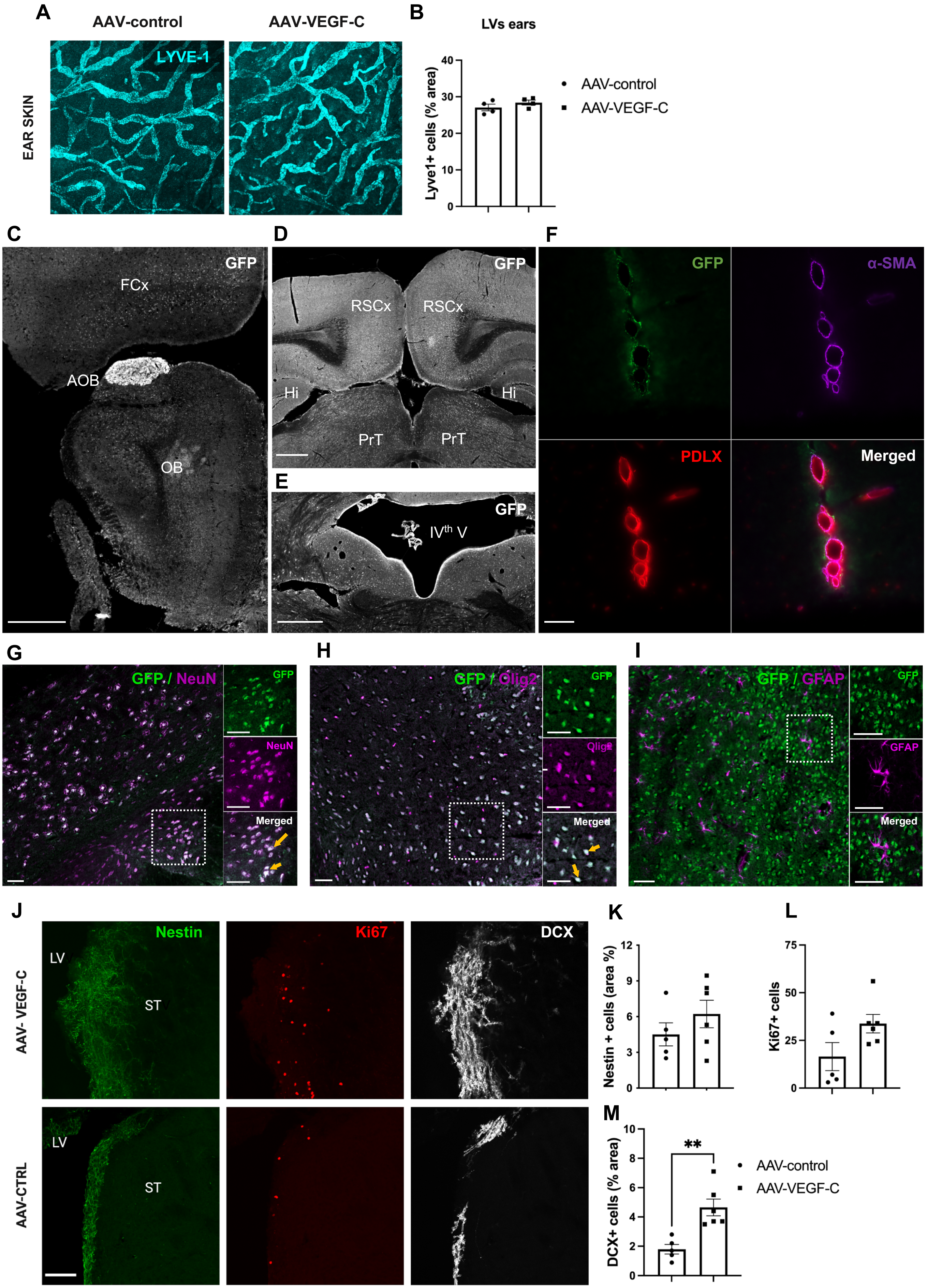

### Supplementary fig 7

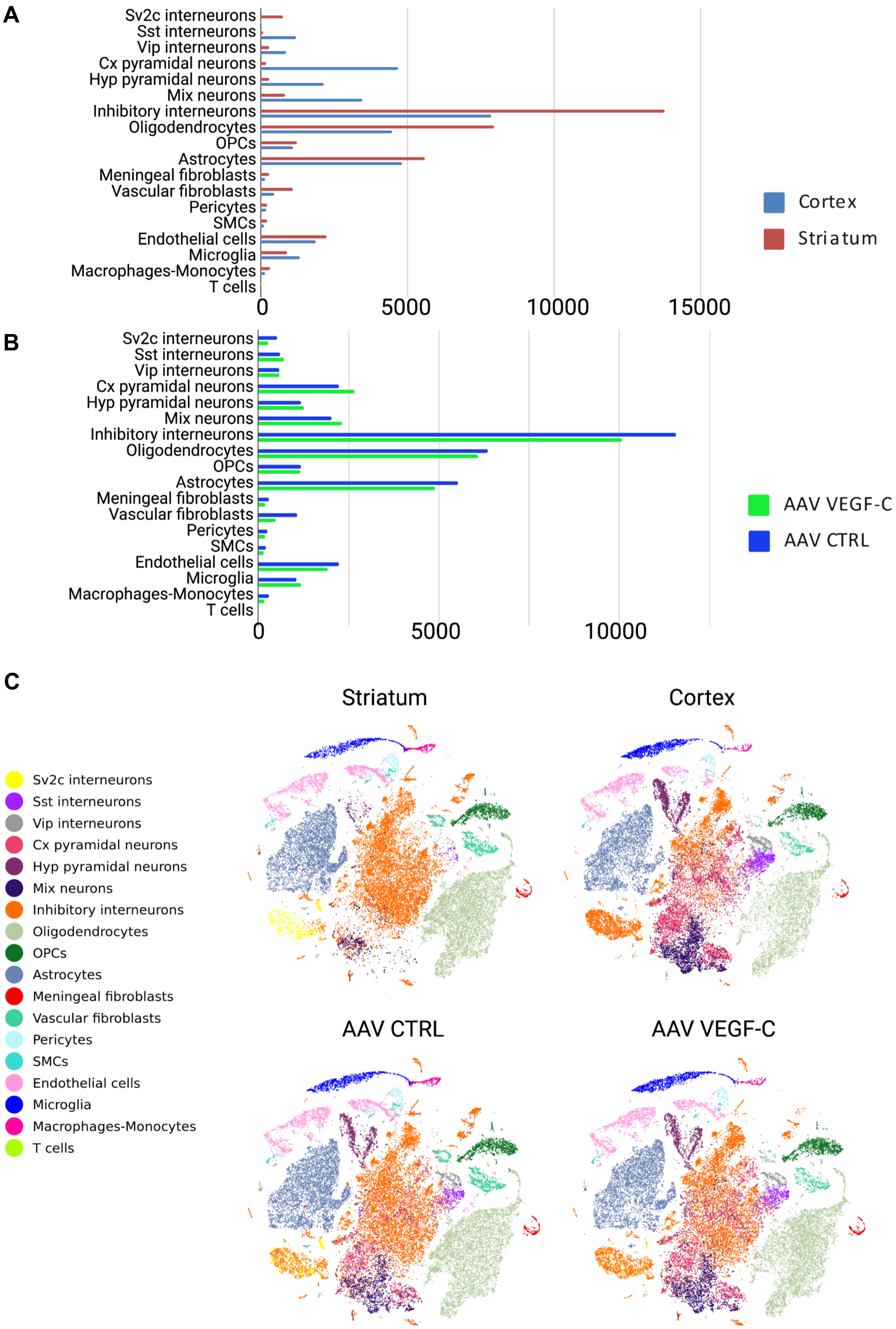

### Supplementary fig 10

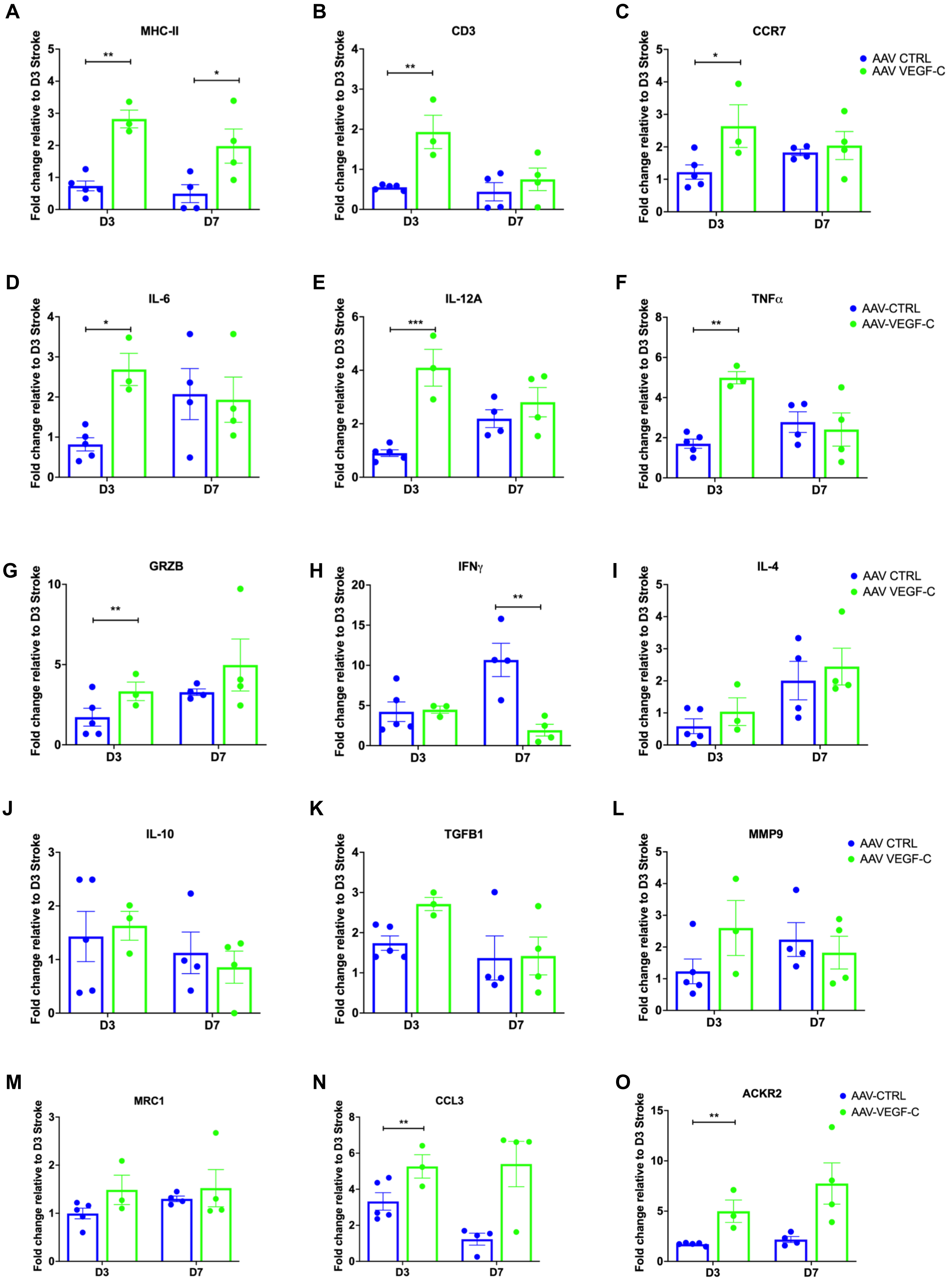
