## Supplementary fig 2 for "VEGF-C promotes brain-derived fluid drainage, confers neuroprotection, and improves stroke outcomes"

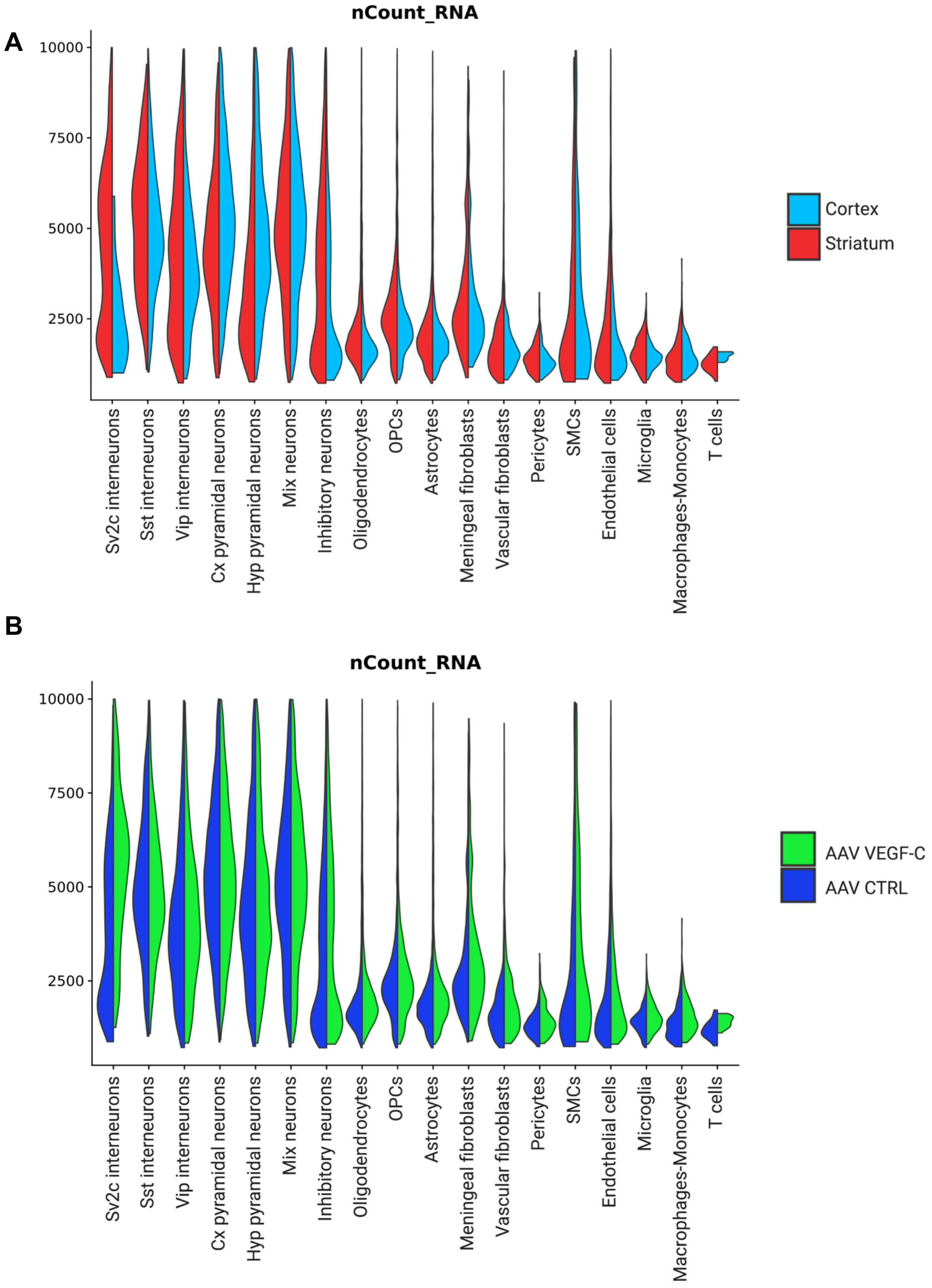


**Supplementary Fig. 2. snRNA-seq data: violin plots representation of transcript number in each cluster**. **(A)** Between regions (Cortex versus Striatum). **(B)** Between conditions (AAV-VEGF-C versus AAV-CTRL) (*n* = 5 mice/group).
