## Supplementary fig 3 for "VEGF-C promotes brain-derived fluid drainage, confers neuroprotection, and improves stroke outcomes"

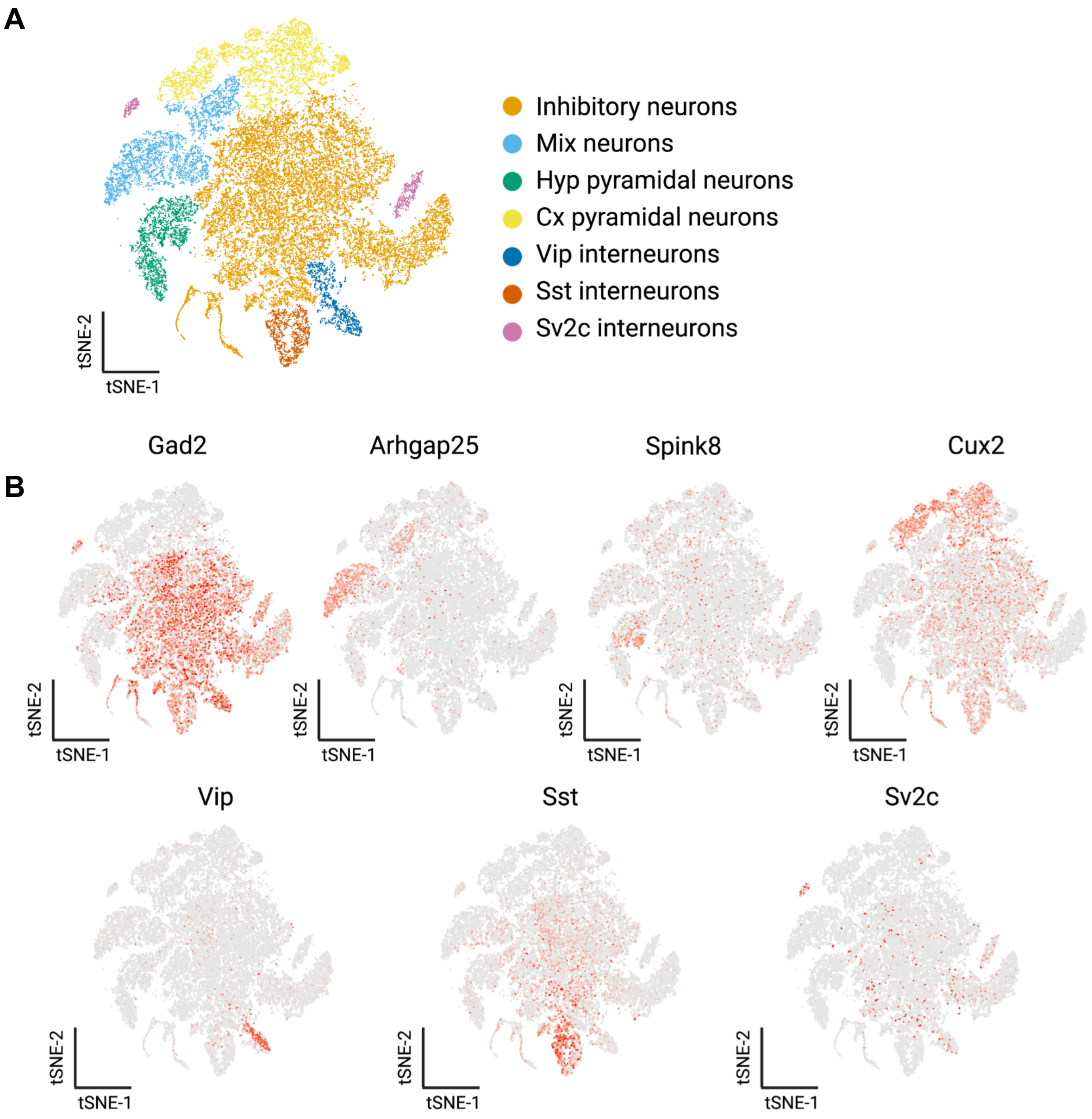


**Supplementary Fig. 3. Sub-clustering of forebrain neuronal cells. (A)** tSNE representation of the neurons after sub-clustering and isolated mapping. **(B)** tSNE representation of marker gene expression in the sub-clusters.
