## Supplementary fig 4 for "VEGF-C promotes brain-derived fluid drainage, confers neuroprotection, and improves stroke outcomes"

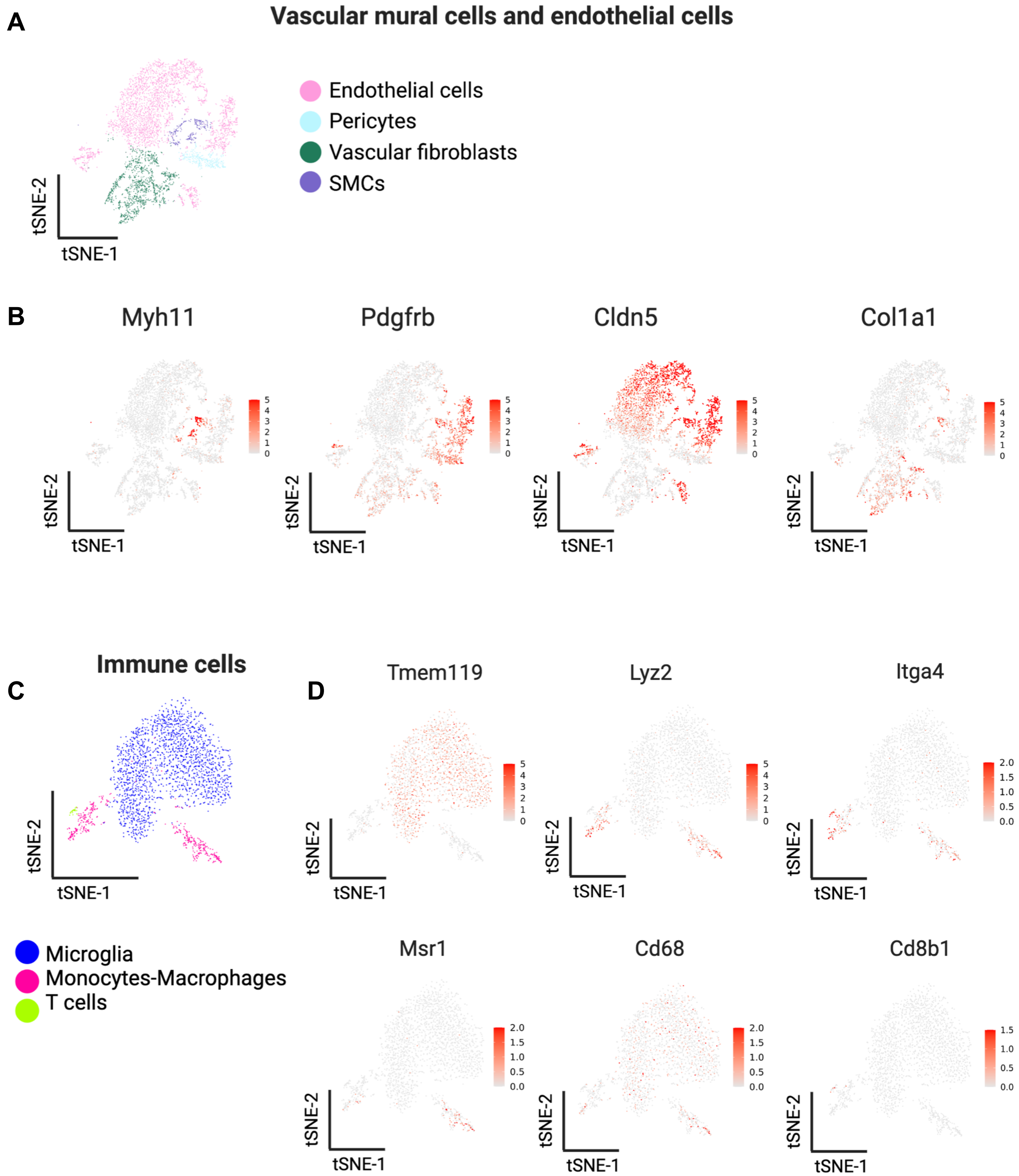


**Supplementary Fig. 4. Sub-clustering of endothelial cells, vascular mural cells, and immune cells.** (**A**) tSNE representation of vascular mural cells and endothelial cells clusters and distribution of relevant marker genes in tSNE representations. (**B**) Endothelial cells (*Cldn5*+), Smooth muscle cells (SMCs) (*Myh11*+); Vascular fibroblast (*Col1a1*+); Pericytes (*Pdgfrb*+ *Myh11*-). tSNE visualization of sub-clusters of immune cells (**C**). Scaled distribution of marker genes of, microglia (*Tmem119*), monocytes-macrophages (*Msr1*, *CD68*) and T-cells (*Cd8b1*) (**D**).
