## Supplementary fig 6 for "VEGF-C promotes brain-derived fluid drainage, confers neuroprotection, and improves stroke outcomes"

**A** Inhibitory interneurons

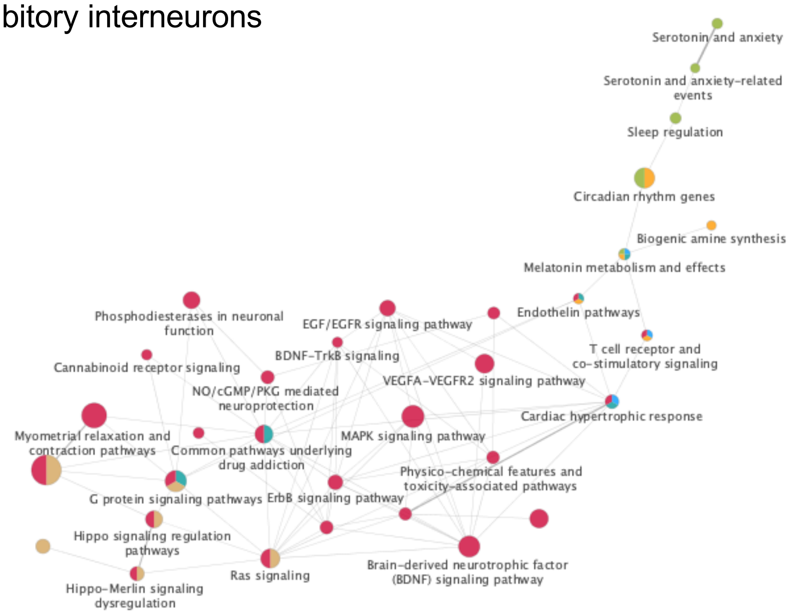


**B** *Sv2c*-interneurons

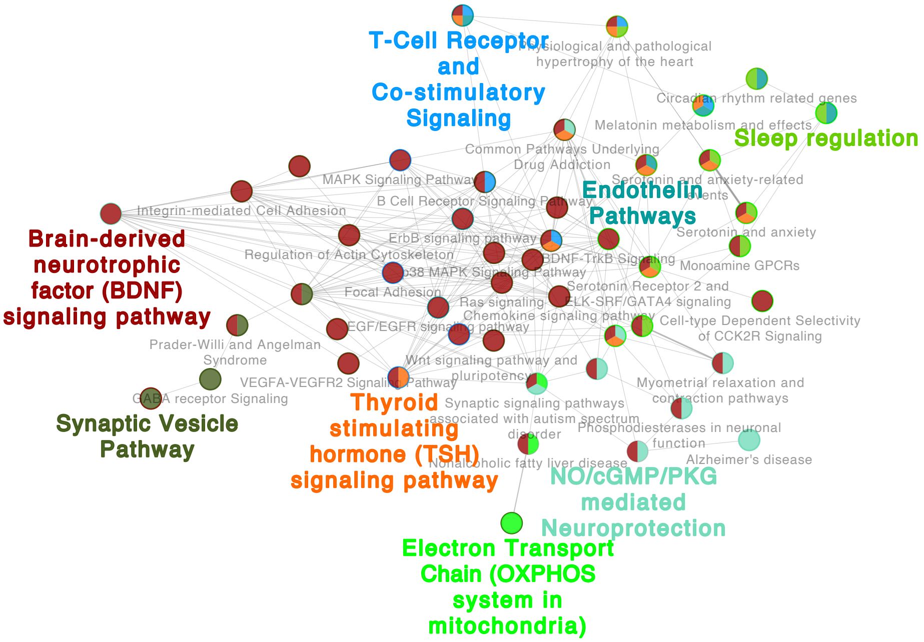


**Supplementary Fig. 6.** Functionally organized network visualized with Cytoscape showing all upregulated pathway interactions in inhibitory neurons (**A**) and Sv2c-interneurons (**B**) following ICM AAV-VEGF-C delivery. Main terms are represented with color. Dot size represents the number of differentially expressed genes in common between pathways.
