## Supplementary fig 7 for "VEGF-C promotes brain-derived fluid drainage, confers neuroprotection, and improves stroke outcomes"

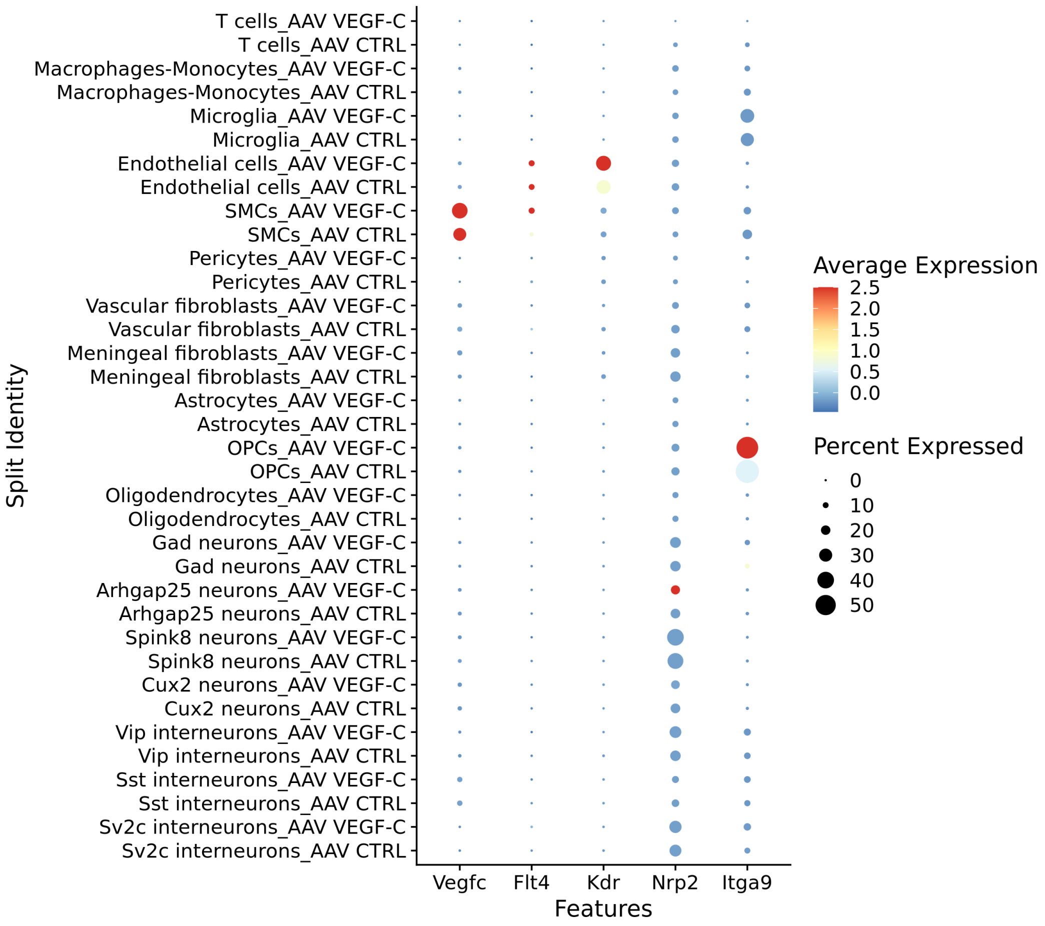


**Supplementary Fig. 7. *Vegfc* and VEGF-C receptors transcript expression**. Dot plot representation of transcript expression among the different clusters using Log normalized and zero centered expression in each cluster (AAV-VEGF-C + AAV-CTRL
