## Supplementary fig 8 for "VEGF-C promotes brain-derived fluid drainage, confers neuroprotection, and improves stroke outcomes"

**Suplementary Fig. 8.**

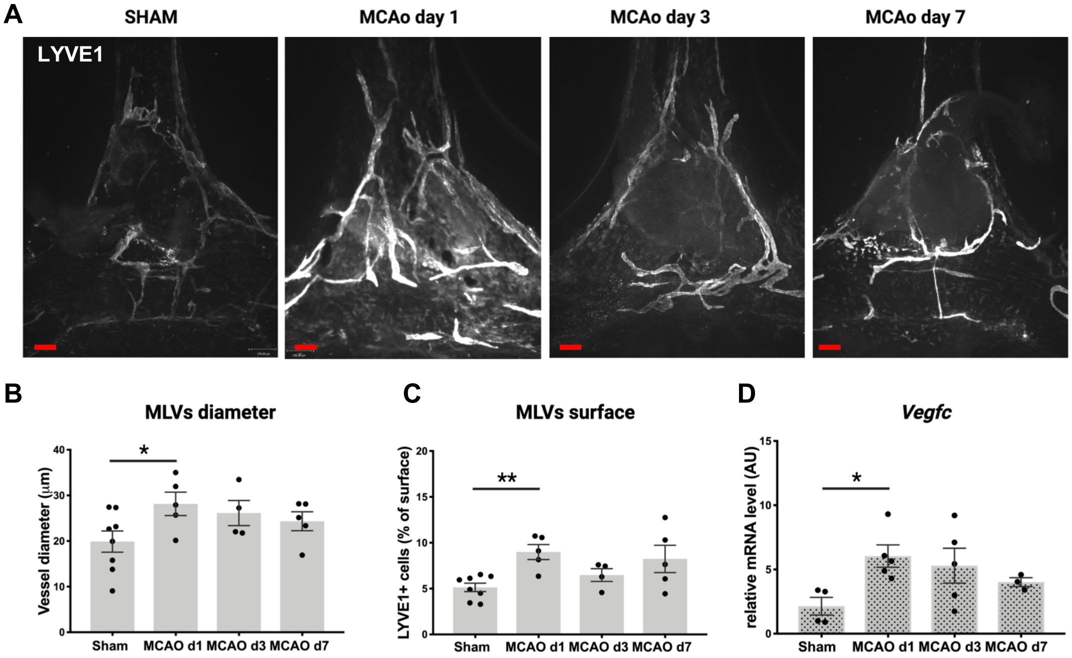


**Supplementary Fig. 8. Transient MCAo induces acute MLV alteration and upregulates *Vegfc* expression. (A)** Illustrative images of Anti-LYVE1-immunolabeled MLVs in the confluence of sinuses in sham mice and at 1, 3 and 7 days after MCAo. Quantification of MLV diameter (**B**) and LVs surface (**C**) at the different time points compared to the sham group ($n$ = 4-8 mice/group, $^{**}P < 0.005$, $^{*}P < 0.05$ Mann-Whitney test). Scale bar: 170 µm. **(D)** Quantification by qPCR of *Vegfc* expression on the right hemisphere (forebrain) ($n$ = 3-5 mice/group $^{*}P < 0.05$ Mann-Whitney test).
