## Supplementary fig 9 for "VEGF-C promotes brain-derived fluid drainage, confers neuroprotection, and improves stroke outcomes"

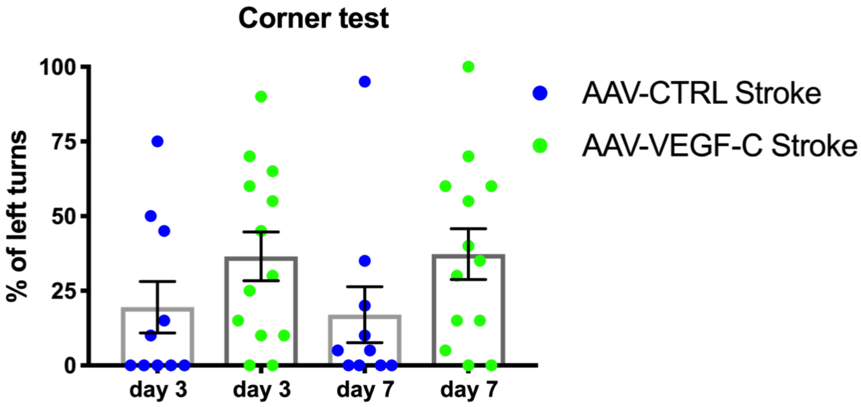


**Supplementary Fig. 9. Corner test evaluation at day 3 and day 7 post tMCAO**. No difference (% percentage) of left turns (impaired side) between AAV-VEGF-C (day 3: 36 ± 8; day 7: 37 ± 8) and AAV-CTRL (day 3: 19 ± 8; day 7: 17 ± 9. *n* = 10-13 mice/group; *P* = 0.21).
